# Assessing risk for butterflies in the context of climate change, demographic uncertainty, and heterogenous data sources

**DOI:** 10.1101/2022.05.22.492972

**Authors:** Matthew L. Forister, Eliza M. Grames, Christopher A. Halsch, Kevin J. Burls, Cas F. Carroll, Katherine L. Bell, Joshua P. Jahner, Taylor Bradford, Jing Zhang, Qian Cong, Nick V. Grishin, Jeffrey Glassberg, Arthur M. Shapiro, Thomas V. Riecke

## Abstract

Ongoing declines in insect populations have led to substantial concern and calls for conservation action. However, even for relatively well-studied groups, like butterflies, information relevant to species-specific status and risk is scattered across field guides, the scientific literature, and agency reports. Consequently, attention and resources have been spent on a miniscule fraction of insect diversity, including a few well-studied butterflies. Here we bring together heterogenous sources of information for 396 butterfly species to provide the first regional assessment of butterflies for the 11 western US states. For 184 species, we use monitoring data to characterize historical and projected trends in population abundance. For another 212 species (for which monitoring data are not available, but other types of information can be collected), we use exposure to climate change, development, geographic range, number of host plants, and other factors to rank species for conservation concern. A phylogenetic signal is apparent, with concentrations of declining and at-risk species in the families Lycaenidae and Hesperiidae. A geographic bias exists in that many species that lack monitoring data occur in more southern states where we expect that impacts of warming and drying trends will be most severe. Legal protection is rare among the taxa with the highest risk values: of the top 100 species, one is listed as threatened under the US Endangered Species Act and one is a candidate for listing. Among the many taxa not currently protected, we highlight a short list of species in decline, including *Vanessa annabella*, *Thorybes mexicanus*, *Euchloe ausonides*, and *Pholisora catullus*. Notably, many of these species have broad geographic ranges, which perhaps highlights a new era of insect conservation in which small or fragmented ranges will not be the only red flags that attract conservation attention.

## INTRODUCTION

Reductions in abundance, contractions in geographic range, extirpation, and extinction have become common features of wild plant and animal populations impacted by the various stressors of the Anthropocene (Dirzo et al. 2014, Turvey and Crees 2019). Effects on individual populations translate into depauperate assemblages of species in remaining natural lands, even those far removed from the most immediate effects of habitat destruction and degradation (McLaughlin et al. 2002, Brook et al. 2008). To the extent that the loss of evolutionary lineages (populations, species and higher taxonomic groups) is a part of life on earth and always has been, the current mass extinction crisis affords ecologists the chance to study extinction as an important earth-system process (Benton 2003). However, the need to maintain functioning natural ecosystems is increasingly generating motivation among scientists, the general public, and governmental bodies to reverse or slow whatever biotic losses might still be addressed (Naeem et al. 2016). Concern for functioning ecosystems has been elevated in recent years by a steady pulse of papers reporting declines in insect abundance and diversity (Eisenhauer et al. 2019, Wagner 2019, Wepprich et al. 2019, van Klink et al. 2020) that have inspired calls for new conservation attention focused on "the little things that run the world" (Wilson 1987, Goulson 2019, Cardoso et al. 2020).

Effective conservation and management actions depend not only on knowing which actions will be effective (Bladon et al. 2022) but also on prioritization of need because resources to support conservation are always limiting. Prioritization in turn depends on the synthesis of multiple lines of information including population monitoring, natural history studies, and geographic surveys. For insects, the taxonomic diversity is so great and the available information is so sparse (Cardoso et al. 2011), that proactive prioritization and conservation informed by diverse data types has rarely been an option. As a consequence, insect conservation has often been motivated largely by fragmentation and small geographic ranges (Samways 2007, Diniz-Filho et al. 2010). Exceptions to that pattern include a few European countries where studies of butterflies and a small number of other insect groups have been sufficiently thorough in terms of natural history and monitoring that researchers have been able to prioritize species for conservation attention in a way that follows the International Union for Conservation of Nature (IUCN) and the Red List framework (Fox et al. 2011, van Swaay et al. 2011, Maes et al. 2012, Bonelli et al. 2018, Franke et al. 2022). That depth of species-specific information for insects is unusual, even for butterflies, and most countries will have a more complex mix of some monitoring or observational data, natural history observations, and expert opinion (New et al. 1995, Edge and Mecenero 2015, Geyle et al. 2021).

Butterflies in the western United States provide an excellent case study for the challenge of conservation prioritization that involves a mixture of heterogenous data types and sources of information. The region does include butterfly monitoring programs, but also expansive areas that are sparsely populated and understudied, in particular the Intermountain West with hundreds of mountain ranges in the nearly 500k square kilometers of the Great Basin Desert. The most temporally-intensive butterfly monitoring program in the western US is the Shapiro transect of ten permanent sites across Northern California that have been monitored biweekly during the flight season for between 35 and 51 years (Shapiro 2022). Many years before the entomological world made a collective pivot to the problem of insect declines (Dirzo et al. 2014, Hallmann et al. 2017), work with the Shapiro data documented shifting spring phenologies (Forister and Shapiro 2003), and the influence of land use and warming temperatures on extensive declines in abundance and species richness (Forister et al. 2010, Casner et al. 2014b). More recently, patterns from the temporally-intensive Shapiro dataset were analyzed in parallel with geographically-extensive monitoring data from the North American Butterfly Association (NABA) and iNaturalist observations across the 11 western states (Forister et al. 2021). That effort quantified a compounding loss of 1.6% fewer butterflies observed per year, consistent with rates of loss from studies in other regions (Wepprich et al. 2019, van Klink et al. 2020), and highlighted the negative influence of warming and drying conditions on butterfly populations in natural areas. However, the species included in Forister et al. (2021) were only those common and widespread enough to be present with sufficient frequency in monitoring databases to allow for inclusion in statistical models. Moreover, an attempt was not made to combine different lines of information into a ranking of species for conservation concern.

Here we address that need by taking a multi-faceted approach to conservation prioritization that utilizes observational data when available (for approximately half the species) and a combination of data types for other species, including natural history traits and quantitative estimates of exposure to climate change and development. We refer to these two groups of species, with and without monitoring data, as the "A group" and "B group" species, respectively. These two groups are treated throughout this paper in distinct but complementary ways, with the dual goal of presenting a unified picture of risk for a regional butterfly fauna, but also with the objective of providing a case study for how species with different amounts of information can be mutually informative when considered together in a conservation context. The different data types studied for both A and B group species are detailed below and are used (1) to produce a quantitative ranking that highlights the taxa most severely declining and most likely to face regional extirpation or extinction in coming decades; and (2) to identify geographic and taxonomic knowledge gaps in our understanding of western butterflies. It is our hope that these results will be used by conservation practitioners and land managers to guide restoration and protection efforts, and will also motivate additional monitoring and the development of new population models that take maximum advantage of heterogenous data types. Throughout this paper, we use the word "risk" (and related terms, like "risk index") in a flexible way that encompasses evidence of past decline, projected declines, and combinations of traits that could predispose species to ongoing and future declines. This flexibility is necessary given the nature of our project encompassing species for which different kinds and quantities of information are available, but in all cases we intend the concept of high risk to flag species that could profitably receive careful attention from ecologists, conservation biologists, and the general public.

## MATERIALS AND METHODS

A schematic overview of our methods is shown in Figure 1, emphasizing the flow of information from external data sources through analyses to the generation of quantitative risk assessment. All parts of the process are discussed in detail here. The order of the sections below follows the order of variables (salmon colored boxes) in Figure 1, from left to right, starting with the process that generates P(persistence) from the NABA data and ending with wing span and host range. After the creation of variables, the subsequent methods sections roughly correspond to the main products (green boxes) in Figure 1, from the creation of ranked lists to risk maps and the visualization of phylogenetic risk.

**FIGURE 1.**
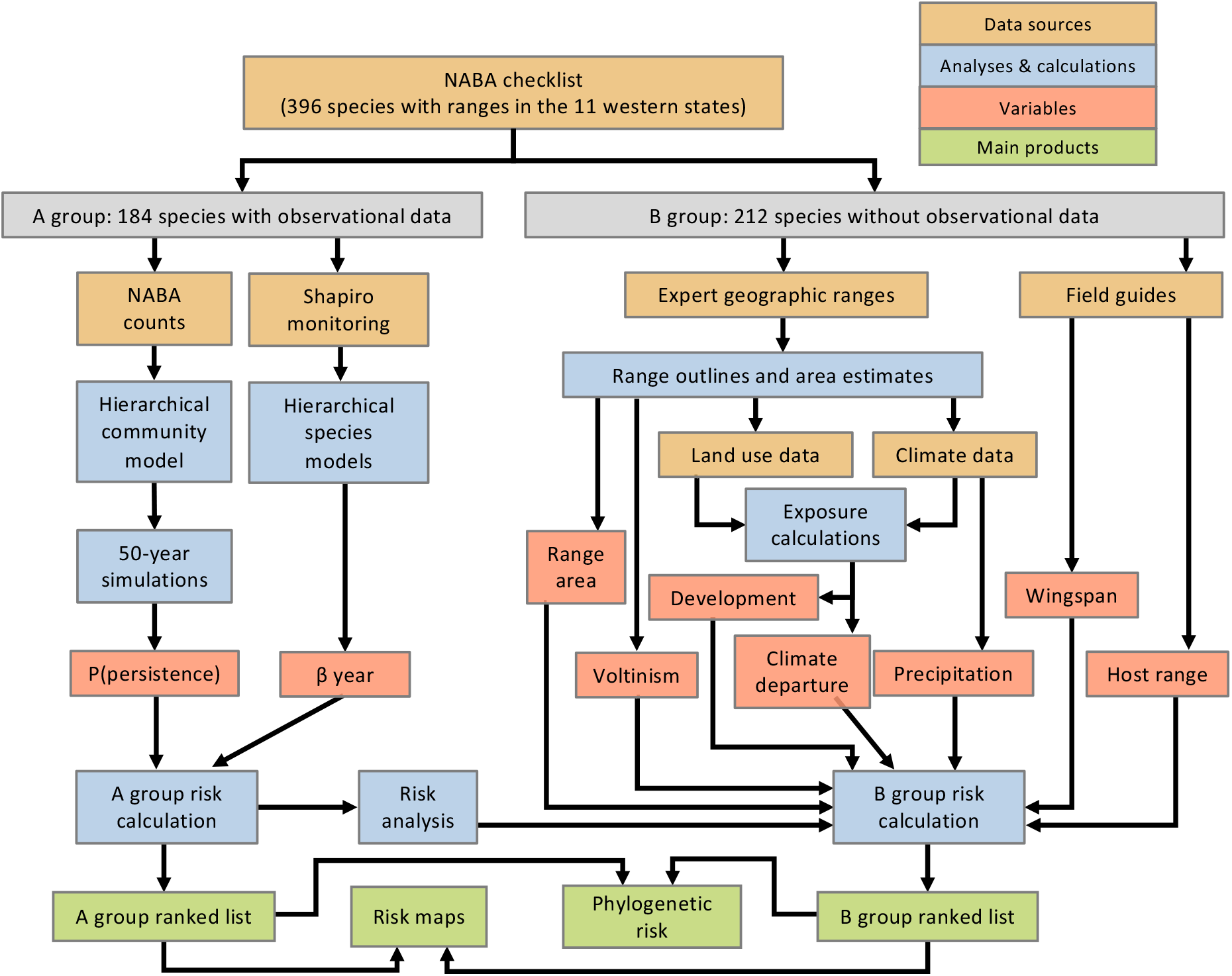
Schematic overview of main inputs, processes and products associated with the generation of risk index values. As noted in the key, data sources are in brown, analyses (and other calculations) are in blue boxes, variables (used in the creation of the risk index) are in red, and the primary products are in green. The central branching path illustrates the division of species into the A and B groups, with monitoring data contributing to the A group risk assessment on the left, and other data types contributing to B group assessment on the right. The 9 variables (in red) are identical to the columns in Figure 2, although labelled slightly different here, especially for the observational variables: "β year" is the year coefficient from analyses of Shapiro data summarizing change through time, and "P(persistence)" is the probability of population persistence from 50-year forecasts. Variables on the right ("range area", "precipitation," etc.) are more self-explanatory. Also note that the expert-derived geographic ranges contribute to the risk index calculations both directly ("range area" and "voltinism") and indirectly as indicated with connecting arrows. Finally, the "Risk analysis" process box (towards the lower left) illustrates the analysis of A group risk that was used to partly inform the weighting scheme for the B group species.

Starting with the 875 taxa on the North American Butterfly Association’s 2nd edition checklist of butterflies occurring north of Mexico (NABA 2018), we retained 396 species with resident (non-vagrant) status in the eleven western states (Washington, Oregon, California, Idaho, Montana, Nevada, Wyoming, Colorado, Utah, New Mexico, and Arizona) based on range maps in Glassberg (2017), and collapsed 18 subspecies from that list into full species. For clarity and in order to facilitate wide use of our results, we also reference a second checklist by Pelham (2022) in places where names differ.

Of the 396 species from the NABA list, 184 were present in monitoring databases (either the Shapiro transect or the NABA count circles) with sufficient frequency to be used in population models. For those species (the A group), our approach is to rank species based on observed and forecast population trajectories. Acknowledging the great uncertainty inherent to insect time series analyses, we present the ranking of A group species in a way that risk associated with other variables (e.g., geographic range size) can be evaluated by the reader.

For the 212 species in the B group (not present in monitoring schemes in sufficient frequency for inclusion in core population models), we have accumulated seven variables that form a composite picture of risk: geographic range, exposure to developed land, exposure to climate change, average (range-wide) precipitation, voltinism (number of generations per year), wingspan, and host range (or "host breadth"). We combine those seven variables into a single risk index as a weighted sum, where the weights are determined in part by our previous work with western butterflies, but also by analyses of the A group (described in detail below). The weighting scheme and other steps in data processing involve informed but partly subjective judgments (see section below on *Calculation of risk index for A and B group species*) with respect to threats to butterflies and natural history traits that predispose butterflies to risk. We have presented all data decisions in a transparent way, so that the reader can judge for themselves the consequences of our methods and decisions, and alternative weights can be assigned by researchers using an online tool (see Appendix S1). In the sections below, we describe first the observational datasets (NABA and Shapiro) and associated analyses, then the seven other variables and how they are combined into composite risk indices and are visualized geographically and in a phylogenetic context.

### Variable creation part 1: North American Butterfly Association (NABA) counts and models

The NABA butterfly count program is a suite of hundreds of individual locations throughout the country that are monitored during midsummer (typically once, but in some cases more than once) by a group of at least four observers recording counts of all individual butterflies seen and identified to species, in a 15-mile (24.14 km) diameter circle. Observations from count circles in the 11 western states encompass different numbers of years at different sites from the 1970s to the present, with the final year in the dataset we examined being 2018 (the data were compiled for analysis in 2019). For the current project, we filtered the observations so that we only included sites that had been monitored for at least ten years, and with the final year being 2017 or 2018 (we did this so as not to generate forecasts for species with a substantial recent gap in observations). More than one monitoring day has been reported per year at a small number of sites, and for those locations we retained only the survey closest to the 4th of July, which is the traditional target date for these censuses. We then excluded any site-by-species combinations in which a species was not present for at least ten years (not necessarily consecutive years). Finally, only species meeting the latter criterion for at least three locations were retained. Those filters resulted in a dataset with 162 species from 44 locations used in the core model and associated population forecasts (we experimented with less stringent filters but found that model performance suffered). For species with less data, we ran a second set of models with lower thresholds, as described after the core model below.

Previous work with the NABA data used hierarchical Bayesian linear Poisson regressions run separately for each species (Forister et al. 2021). Here we advance that approach using a single, multi-species model that shares information about heterogeneity in the observation process across species observed at each site (Riecke et al. 2021). The components of the model (each described in turn below) include an observation sub-model, an abundance sub-model, and a forecast or simulation process that projects occupancy (the fraction of sites with non-zero presence by species) for various intervals of years in the future.

For the observational component, we modeled the counts of individual butterflies (*y*) using a Poisson distribution given the expected count of each species at each location during each year (*μ*_t,l,s_), where *t*, *l*, and *s* identify the year, location, and species respectively:

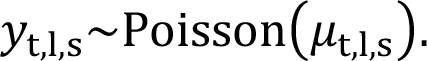

We modeled the expected count (*μ*_t,l,s_) as a function of an abundance index (*N*_t,l,s_), year- and site-specific survey effort (*β*), and a year- and location-specific random effect (*δ*_t,l_) shared among species:

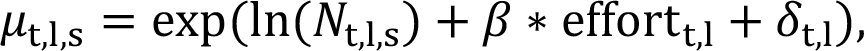

with a vague prior for the effect of survey effort:

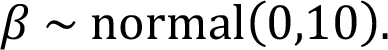

The empirical variable for effort is the z-standardized total hours searched by all survey groups at a site on a day. After accounting for the effect of survey effort, we modeled additional variation in detection probability for each survey or monitoring day as a random effect shared among species. This random effect can be thought of as the combined effects of survey-specific variation in detection due to processes such as variation in observer experience and local weather conditions (Riecke et al. 2021):

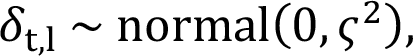

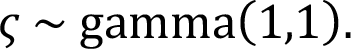

For the abundance sub-model, we assigned priors for initial population abundance indices for each species at their first encounter (*f*_site_*i*_,species_*i*__) at a study site as a function of initial survey effort and the initial count:

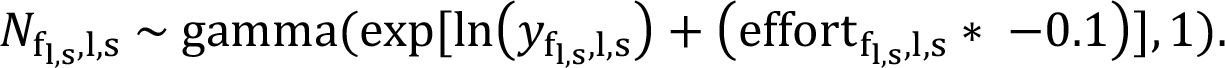

We modeled changes in population size (N) from one year to the next for each species at each site as a function of year (t), location (l), and a species(*s*)-specific population growth rate (λ):

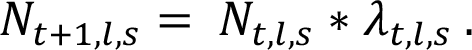

Variation in population growth rate was in turn modeled as a function of a species-specific mean population growth rate (γ_s_), and species-specific random variance in population growth rate:

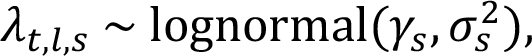

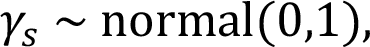

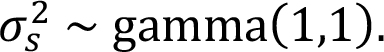

Finally, for each species at each location, we projected the abundance index into the future using Monte Carlo simulation from the posterior distributions of species-specific population growth rate (*λ_t,l,s_*), and species-specific population growth rate variance 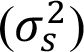:

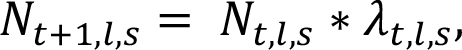

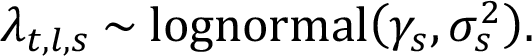

We defined local ‘extirpations’ as locations at which the expected count of a species given mean effort was less than 0.1 individuals, and calculated extirpation probability for each species at 10, 20, and 50 years into the future. Thus, one minus the extirpation probability is the probability of population persistence, and it is that value (probability of persistence) for each species from the core NABA model that moves forward (represented by 1k samples from the final year of the simulations) into the calculation of the risk index for the A group species.

The above model and 50-year projections were used for 162 species in the A group with a sufficient number of observations (passing the filters described above); note that the total number of A group species is 184, not 162, because another 22 species are in the A group as a consequence of their presence only in the Shapiro dataset (107 A group species are present in both datasets). For a different set of 105 species in the B group, we used a less complex model than the one described above. These species (a subset of the B group) were present in the NABA dataset, but with too few observations (a median presence of 2 sites per species) to be included in the core NABA model. However, in the interest of presenting maximal information on all species and using all available data, we estimated trends through time for this subset of the B group; the results are reported but not incorporated into the risk index calculation for these species. In this model, the counts (*y*) were also modeled with a Poisson distribution given the expected count for each location and year (*μ*_t,l_), where *t* is the year and *l* is the location:

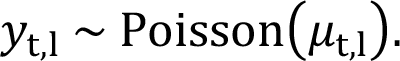

The expected count (*μ*_t,l_) was then modeled as a linear function of a site-specific intercept (*α*_1_), a site-specific (s) year effect (*β*_1_), and site-specific effect of effort (*β*_2_):

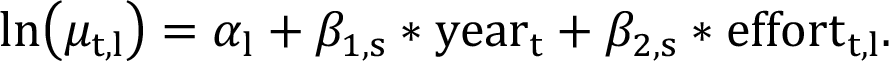

The intercept and both beta coefficients were drawn from normal priors, with the normal truncated at zero to be positive for effort (*β*_2_); the means and variances of those distributions were in turn drawn from hyperpriors (thus estimating effects across sites) with means drawn from normal distributions (with mean of zero and variance of 100) and variances drawn from gamma(1,1) as in the core model above. For 35 species present at only a single site, the model was run without the hierarchical (across sites) structure. The output of these secondary models (for the 105 species) was retained as a directional probability (the fraction of the posterior distribution above zero for species with a positive year coefficient, and below zero for species with a negative year coefficient).

All Bayesian models were implemented using JAGS (version 4.3) and the `jagsUI` package (Kellner 2017) in R (R Core Team 2020). The core model (for A group species) was run with three chains for 500k iterations, with a 250k iteration burn-in. The secondary models (for the 105 B group species with some presence in the NABA data) were run with two chains for 2k steps and a 1k burn-in. Model diagnostics included inspection of plots of chain histories (all chains converged; 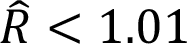), and effective samples sizes.

### Variable creation part 1b: Phenology

A potential issue with once-per-year sampling, as in the NABA data, is sensitivity to phenological shifts. For example, earlier spring emergence associated with a warming climate could cause a mid-summer census date to fall after a population peak, and thus appear as a reduction in population density. This might be especially relevant for species with fewer generations per year (univoltine or bivoltine species) where average flight date tends to be correlated with emergence date (Wilson et al. 2007), but could be relevant for any species depending on the timing of peak population productivity. On the other hand, we have found that earlier emergence for multivoltine butterflies in low elevation California tends to be associated with elevated numbers throughout the summer as populations have more time for growth (Forister et al. 2018). To consider these possibilities here, we developed an alternative version of our core Bayesian model (described above) in which sampling date is included as a linear covariate with effects estimated separately for univoltine and multivoltine species. Sampling date in the NABA dataset varies among sites and years, and has been advancing at the rate of a little over one day per decade when estimated across sites (Appendix S1: Figure S1). Results from that model, including effects of sampling date on the different voltinism groups, were nearly indistinguishable from the results of our primary model (see Appendix S1: Figure S1 for this and other relevant results), thus in the interest of parsimony we do not include the phenological term in our core model and results presented throughout the rest of this paper.

We also used the more temporally intensive Shapiro dataset, with biweekly samples (described in detail in the next section), to further explore potential biases with once-per-year census data (as in the NABA dataset). These analyses focused on the years since 1999 at the five low elevation Shapiro sites, for which counts of individuals are available, as opposed to presence and absence observations that are available for earlier decades at all sites. Using species with at least 10 years of data, we calculated the following indices for each species at each site, using generalized linear models with Poisson error and a log link function: (1) rate of change in the date of first flight, estimated as the year coefficient from models predicting the date of first observation in each year; (2) rate of change in total annual abundance (summed across all visits), as the year coefficient from models that included (in addition to year) the number of sampling days as a covariate for effort; (3) rate of change similar to the previous, but instead of total annual abundance the dependent variable was the count of individuals on a randomly selected day that a given species was observed in a given year; day of the year was included as a covariate for phenology. For the third index, change in abundance based on simulated once-per-year sampling, the single day in each year was restricted to the window between June 15 and August 15 to correspond to variation in mid-summer count circles hosted by NABA. With those three indices, representing change in phenology, change in annual abundance, and change in abundance on a single census day per year, we used simple Pearson correlations to ask if species that have emerged earlier since 1999 would appear to be in decline based on the once-per-year sampling, as predicted by the idea of a missed peak of abundance (discussed above). We also asked if any trends in abundance detected with the simulated single census point data would align with change over time inferred from analysis of counts of abundance summed over all visits in a year.

### Variable creation part 2: Shapiro transect data and models

Ten long-term study sites across northern California have been monitored for between 35 and 51 years (depending on the site), with the presence of all butterflies noted along fixed routes every two weeks during the flight season. Data used here were compiled in 2021, including observations through 2020; earlier years were truncated so the dataset starts at 1985, except for three sites where data collection began in 1988. Species by site combinations of at least eight years were retained for analyses of 133 species. Additional details on sites, butterflies and field methods have been described elsewhere (Forister et al. 2010, Halsch et al. 2021, Shapiro 2022). In brief, data from the Shapiro sites have been analyzed using hierarchical Bayesian linear models in which the response variable (the number of days a species is observed in a year) is modeled as a binomial process, with a beta coefficient from the year term in the linear model representing change through time in the probability that a species is observed (Nice et al. 2014, Halsch et al. 2021). Here we use the version of this model and implementation as described in Forister et al. (2021) in which the model was run separately for each species and beta coefficients for years are estimated within and across sites; the higher level coefficients (across sites) are used as indices of population change for each species across the northern California sites. As with the NABA models, model diagnostics included inspection of convergence and effective sample sizes. For downstream analyses (the creation of the risk index for A group species), 1k samples were retained from the posterior distributions of the year coefficients estimated across sites for each species. For two species, *Lycaena rubidus* and *Agraulis vanillae*, the year coefficients were extreme outliers (in the negative and positive direction, respectively) and were not used in the creation of the risk index values (described below) but we do include those coefficients in visual summaries of patterns across species.

The year coefficients from this modeling approach have been shown to be effective indices of change in total abundance as reflected in total counts of individuals which are available from a subset of years and sites (Casner et al. 2014a). Unlike the main NABA model, described in the previous section, we have not taken a forecasting approach with the Shapiro data. The two datasets have different strengths and weaknesses. The strengths of the Shapiro data are intensity and consistency of observation, which lend precision to estimates of species-specific change through time. In contrast, the NABA observations are only once per year, but because they are counts of individuals for all years they can be used to estimate population growth rates (see model description above) which can in turn be used to forecast population occupancy.

### Variable creation parts 3 and 4: Geographic ranges and voltinism

Expert-derived range estimates from Glassberg (2017, 2018) were generated from Keyhole Markup Language (.kml) files for each species. The range of each species was separated by voltinism (the number of generations per year in different portions of the range), with spatial polygons retained separately for uni-, bi-, and multivoltine regions. Quantitative area estimates were then derived for each species within the US and Mexico using the area function in the R package `raster` v3.5-11(Hijmans et al. 2021), which estimates area based on the size of raster cells. For species with ranges spanning from the 48 states into Canada and Alaska, we scanned range maps from Scott (1986) and quantified those northern range portions with the software ImageJ (Collins 2007). These area estimates were added to the values derived from the other sources (Glassberg 2017, 2018). In general, area estimates become biased closer to the poles, but our use of range estimates is relative, and we expect the imprecision to have little effect on results.

In addition to estimating the total area of geographic ranges, the outlines of the expert-derived ranges were used for multiple purposes, including calculating the fraction of each range that is univoltine. For simplicity, we focus on a univoltine vs bivoltine plus multivoltine comparison, rather than considering bivoltinism as a distinct category. The outlines of the expert-derived ranges were also used to calculate exposure to land use and climatic variables (as described in the next section). Note that these other lines of geographic information, from voltinism to exposure to development and climate change, are focused on the 11 states of the western US and do not consider, for example, climate change in other regions.

### Variable creation parts 5 through 7: Land use, climate, and climate change

Previous work with butterflies in our region has revealed effects of land use and climate change that are complex, potentially interacting, and dependent on both the species involved and the landscape context (Casner et al. 2014b, Forister et al. 2018, Halsch et al. 2021). Summarizing exposure to land use and climate change is not a simple task, but we have taken the relatively straightforward option of using the range outline (described in previous section) to quantify these stressors within the range of each species. Note that this differs from the use of point locations to quantify proximity to, for example, urban development (Jamwal et al. 2021). The range-outline approach is a better fit for our goals simply because all species have the same starting data (the expert-derived ranges), which would not be true of 396 species using available point-occurrence records in, for example, iNaturalist. For highly mobile animals, like butterflies, the range-outline method has another advantage in that we do not have to assume that point locations of observations represent the only or most relevant habitats.

To quantify land use change, we reclassified the 2020 Cropland Data Layer (USDA 2020) into land cover types of agriculture, development, or natural and semi-natural habitats using the associated Cropland Data Layer scheme; all crops were classified as agriculture, development of any intensity level as development, and remaining land cover types (including pastureland) as natural or semi-natural habitat. For each species, we used the spatial polygon generated from the range map to clip the rasterized land cover types and calculated the proportion that was agriculture or development. Because of the form in which we acquired the spatial data, this process was done separately for regions of different voltinism, and these were summed to a single value for each species (see Appendix S1: Figure S2 for examples of range-wide exposure to land use).

To estimate climate change exposure, we used TerraClimate data for minimum temperature, maximum temperature, and precipitation (Abatzoglou et al. 2018), which we resampled from ∼4km^2^ spatial resolution to ∼40km^2^ for computational efficiency. Using multivariate Mahalanobis distance as a measure of departure (Farber and Kadmon 2003, Abatzoglou et al. 2020), we calculated departure from baseline conditions (1958-1987) for the most recent thirty years (1991-2020) for each cell. To estimate exposure to climate change, we calculated rate of change in departure over time using Theil-Sen slopes (Theil 1950, Sen 1968) which estimate the median slope between each pairwise set of observations and are relatively robust to outliers near the start or end of a series. We generated a raster of these trends in departures for the eleven western states. For each species, we then clipped the climate departure raster layer using the species range maps as spatial polygons and calculated the mean climate change exposure across that portion of the range (as with land use, this was done separately by voltinism, but then added for a single value per species for further analyses; see Appendix S1: Figure S2 for examples). We also calculated 30-year climate normals (1991-2020) for minimum temperature, maximum temperature, and precipitation annually and within each season across the entire range for each species. Among those three variables, precipitation was recently found to be predictive of changes in butterfly abundance across the west (Forister et al. 2021), thus it was used as a static description of climate for inclusion in the composite risk index (described below).

### Variable creation parts 8 and 9: Wingspan and host range

Among the many morphological and natural history traits that could be informative of status and risk, body size and ecological specialization are widely studied, and thus relevant data are available for many species. More narrow diets are often associated with greater sensitivity to habitat loss and other disturbance (Hughes et al. 2000), and dispersal ability is a key determinant of metapopulation resilience in the face of fragmentation or other stressors. Wingspan has been shown to be a proxy for dispersal ability in butterflies (Sekar 2012), and the values derived here were taken primarily from Opler (1999), and also from Warren et al. (2013). Similarly with diet breadth (or host range), we used a single source for the vast majority of species (Scott 1986), supplemented with Brock and Kaufman (2006) and Lotts et al. (2007).

We gathered both the number of plant genera and plant families reported as caterpillar hosts for each species, and then calculated a combined index of diet breadth as the number of taxonomic families plus the natural log of the number of genera. This calculation of taxonomic diet breadth puts most weight on the number of families but allows for some influence of the number of genera eaten. For example: a species that uses hosts in two genera in two families would have a diet breadth of 2.69 (2 + ln(2)), while a species that uses plants in three genera in two families would have a diet breadth of 3.10 (2 + ln(3)). We did not attempt to gather species-level host records, for which too much data would be missing or unreliable.

### Products part 1a: Transformations prior to risk index calculation

In total, we compiled nine variables that contribute to the prioritization of A and B group species in different ways: (1) 50-year occupancy projections (probabilities of population persistence) based on NABA data; (2) historical rates of change from the Shapiro data; (3) geographic range based on expert assessment; (4) exposure to agricultural and other developed lands; (5) exposure to climate change; (6) average precipitation throughout the range; (7) the fraction of the range with one generation per year; (8) wingspan; and (9) an index of dietary specialization or host range (Figure 1). Prior to their use in assigning a risk value to each species (discussed in the next section), each variable was subjected to a specific set of transformations that resulted in a variable with a range of 0 to 1 where larger values represent greater risk. Depending on the nature of the variable (when larger values do or do not naturally represent higher risk), the transformations included a change of sign, and (for all variables) standardization between 0 and 1 (by dividing by the largest value). In some cases, for highly skewed variables, a natural log transformation was applied as the first step, and all transformations and scaling steps are illustrated in Appendix S1: Figure S3.

For visualization of the transformed and scaled variables and for comparison among species, we divided the distributions (Appendix S1: Figure S3) into quantiles and assigned circles of different sizes to the different intervals, with larger circles indicating larger values and greater assumed risk. For most of the variables, we found that the following breakpoints provided a useful assignment of circles for visualization: 0.15, 0.5, and 0.85; in other words, the interval from 0 to 0.15 was assigned the smallest circle (the least risk), from 0.15 to 0.5 the next largest, etc. Breakpoints differed for some of the more skewed variables (e.g., host range), but the results are interpreted in the same way (larger circles represent larger assumed risk).

### Products part 1b: Calculation of risk index for A and B group species

Here we discuss how the variables described in the sections above are combined into a weighted sum that becomes the risk index for each of the 396 species. This process happens in parallel for the A and B group species, but these processes are not entirely disconnected, as the monitoring-based risk index for the A group is studied in relationship to the other variables (geographic range, voltinism, etc.) for that group, and the lessons learned from that analysis inform the structure of the weighted sum for the B group species.

The A group taxa are those species for which data were available from at least one of the monitoring programs, the Shapiro transect or the NABA network of count circles. For these species, we calculated a weighted sum based on those two lines of information with weights split evenly between them: 50% NABA and 50% Shapiro. Thus, a species with the most severe declining values for each dataset would receive a composite risk score of 1. To incorporate uncertainty retained from Bayesian analyses of the NABA and Shapiro data, the composite risk index was recalculated 1k times using 1k samples of the relevant posterior distributions; we then calculated a mean and 85% highest density interval of risk for each species. Alternative weighting schemes among all variables (including the two monitoring variables) can be explored using an interactive, online tool; see Appendix S1.

The B group species are those lacking monitoring data. Thus, we used a composite of the other seven variables to estimate risk. We experimented with a number of weighting schemes for those seven variables and settled on an approach that was partly influenced by previous research (e.g., Forister et al. 2021) but also informed by an additional analysis of the species in the monitoring data. Specifically (for that additional analysis), we took the composite risk index for the A group species (based on NABA and Shapiro data) and used a linear regression model to determine which of the other seven variables were most predictive of that risk index (following general protocols with other Bayesian models as described above). The exact weighting scheme for B group species (influenced partly by results of the analysis of the A group) is described fully in results below. Clearly many schemes are possible for a weighted sum of seven variables, and we report correlations among outcomes from different schemes. Finally, many of the B group species had some data from the NABA dataset that were not sufficient for inclusion in our main model and occupancy forecasts. For those species, we ran a less complex model (described above as the secondary set of NABA models) and report the results along with other B group results, but we do not incorporate those values into the B group risk index to maintain consistency in risk index calculations.

The calculation of the risk index for both the A and B groups relied on a complete data matrix. For most of the variables used for the B group, there were no missing values, specifically for all of the variables deriving in part from the expert geographic ranges: range area, voltinism, precipitation, development, and climate departure (Figure 1). A few species lacked data for host range, and these we filled with interpolation of the median value calculated across all species.

Similarly, median interpolation was used with the observational data and the A group species. In other words, a species without sufficient NABA observations for analysis was given the median value associated with that variable (across other species in that dataset) prior to the calculation of the risk index.

### Products part 2 and 3: Geographic and Phylogenetic visualization of risk

Finally, we asked how the composite risk indices were distributed across the landscape and across the phylogeny of western butterflies. From a spatial perspective, we calculated both the cumulative estimated occurrence of at-risk species (separately for each cell in a raster covering the extent of the eleven western states) and average risk among species present in a cell. We did this separately for the A and B group species, and we restricted analyses to only species with higher risk values by subsetting to the upper 75th quantile of risk values separately for each list (A and B). Within those higher-risk groups, we converted each species range map from a spatial polygon to a raster layer where values within the range were set to 1 and values outside the range to 0. We summed these values across all rasters to produce a new raster of cumulative estimated species occurrence (also referred to simply as "cumulative occurrence"). To calculate mean risk for each cell, we divided the cumulative risk index raster by the cumulative occurrence raster.

For the evolutionary perspective, we used the phylogeny from Zhang et al. (2019) for all 845 butterfly species from the United States and Canada. Briefly, this tree was based on 756 universal single-copy orthologs we identified from 36 reference genomes using `OrthoMCL` (Li et al. 2003). Sequences of these orthologs were aligned using both local (BLAST [Altschul et al. 1997]) and global (MAFFT [Katoh et al. 2002]) alignment methods, and only positions that were consistently aligned by both methods were used. Sequences of non-reference species were derived by mapping the Illumina reads to the exon sequences of the reference species and performing reference-guided assembly. Multiple sequence alignments (MSA) of different orthologs were concatenated to a single MSA. This MSA was partitioned by codon position and used to build a tree by `IQ-TREE` (version 1.6.12) (Nguyen et al. 2015) with the most suited evolutionary model automatically found by `IQ-TREE`.

The phylogeny was imported as a time-calibrated .tre file into R and pruned to our focal western butterflies (the combined A and B group lists). The package `ggtree` (Yu et al. 2017) was used to plot a phylogeny with tips labeled by risk categories assigned based on the quantiles of the risk distributions separately for the A and B group species. Specifically, species in the upper 90th quantile were labeled as "high risk," species between the 75th and 90th quantiles were labeled as "medium risk", and species below the 75th were "low risk." Finally, the phylosig() function from `phytools` (Revell 2012) was used to calculate lambda and K (with 1000 simulations for the permutation test) as measures of phylogenetic signal for the continuous risk index across all species, which in this context is informative with respect to the extent to which closely related species share similar levels of risk.

## RESULTS

We calculated an index of risk for 396 species, which includes two groups: 184 species in the A group with extensive monitoring data, and 212 species in the B group without observational data (or without enough to be used in our primary population models). The B group species tend to have smaller geographic ranges (Appendix S1: Figure S4), which in part explains their reduced presence (just by geographic chance) in monitoring programs, but the two groups differ in other ways. The B group species have slightly lower exposure to development (Appendix S1: Figure S4) and moderately higher exposure to climate change (Appendix S1: Figure S4). The higher climate change exposure is explained in part by the greater presence of more southern species in the B group, as seen by latitudinal midpoints (Appendix S1: Figure S4) and qualitative characterization of range (Appendix S1: Figure S4).

For the A group species, we modeled historical and projected population trajectories using different sources of observational data. Consistent with previous work with NABA data, our new model with shared (across-species) observation heterogeneity found that a majority of species (71%) had annual population growth rates below replacement (Appendix S1: Figure S5). We used those estimated annual growth rates and the most recent year of observed counts to simulate 50 years into the future. The median fraction of extant locations (or probability of local persistence) per species at 50 years was 0.60, and that fraction was positively related to historical population growth rates (Appendix S1: Figure S5). Results from analyses of Shapiro data also find that a majority of species exhibited downward trends through time of varying magnitude (84.5% of species had negative year coefficients).

Our risk index and ranking was based on a combination of evidence from the NABA and Shapiro datasets, but note that the A group species are shown in Figure 2 with risk information associated with the other seven variables (geographic range, exposure to development, etc.), even though the actual ranking of the A group is based solely on the observational data. We present the information in this way because we acknowledge the imperfect geographic coverage of monitoring programs and the inherent uncertainty in population models. Thus, the reader or conservation practitioner can easily see if two species with similar risk values in the A group (based on NABA and Shapiro) potentially have similar risk based on other variables like range size.

**FIGURE 2.**
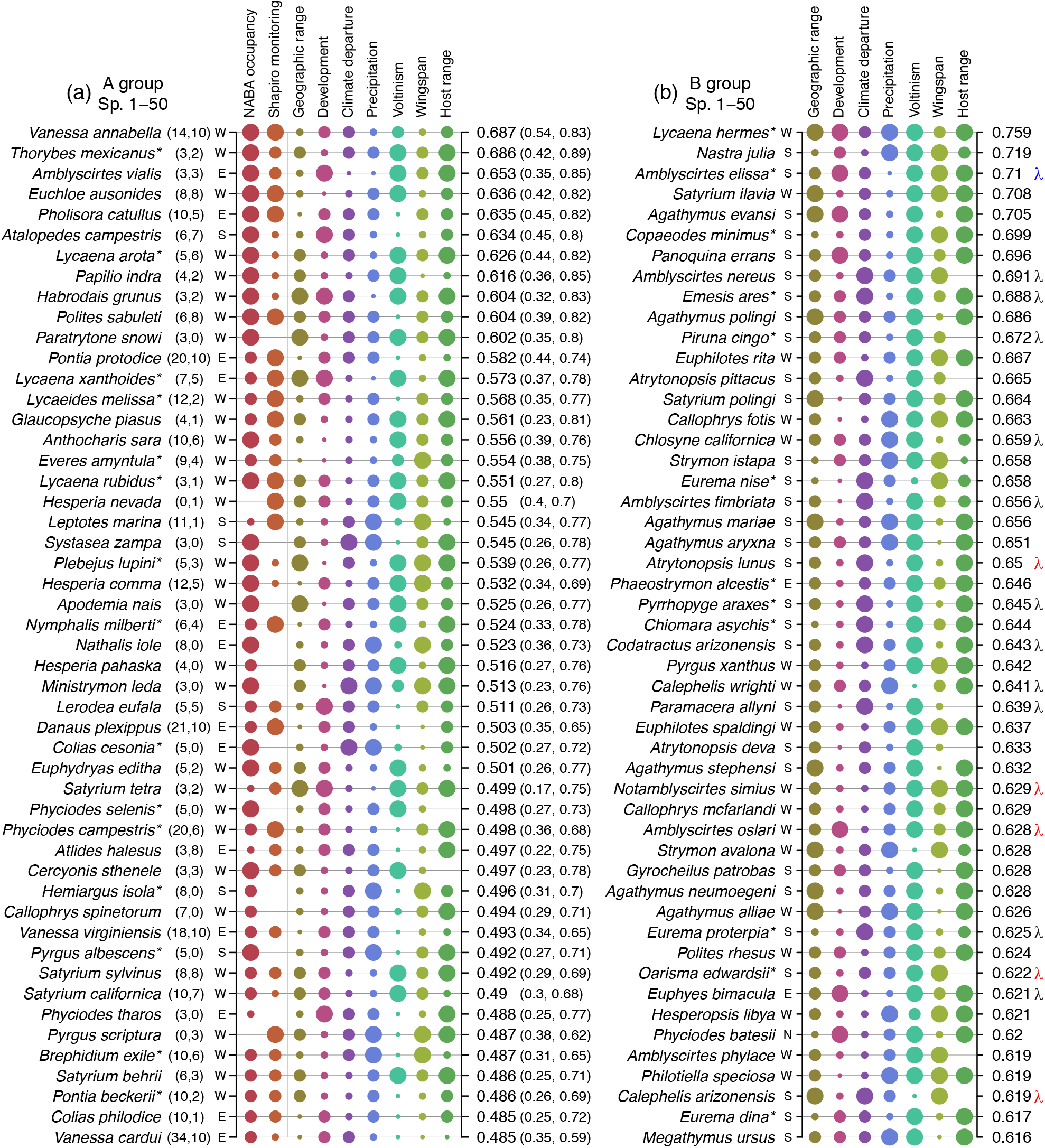
The top 50 species with the highest risk values in the A group (on the left) and the B group (on the right). The two panels have some features in common, and some unique elements. In common they both show the extent to which different variables are associated with higher or lower risk for each species: a large circle under NABA occupancy, for example, marks a species that we infer as being at risk because of low forecast occupancy (probability of population persistence) across currently-extant locations; similarly, a large circle under development indicates a species at risk because of high exposure to developed lands, and a large circle under geographic range indicates corresponding risk associated with a relatively small range. The sizes of the circles were assigned separately within the two lists, A and B group species, and thus indicate relative differences within those lists. Although all variables are shown for comparison, the overall risk ranking for the A group species is based solely on the first two variables (NABA occupancy and Shapiro monitoring, to the left of the vertical gray line), while the ranking for the B group species is based entirely on the other seven variables (see main text for details, and Figure 1). Both panels also have in common the quantitative risk values shown to the right (e.g., the risk index for *Vanessa annabella* in panel A is 0.687); note that the risk values for the A group species include 85% credible intervals (in parentheses), encompassing uncertainty derived from Bayesian analyses of both NABA and Shapiro data. The capital letters (N, S, E and W) running down the left side of each panel are qualitative biogeographical descriptions indicating where the mass of a geographic range lies relative to the western US (N and S indicate species found primarily north and south of the US borders with Canada and Mexico, respectively; W indicates species with the majority of their range in our focal region; and E indicates species found either mostly in the eastern US or with a transcontinental distribution), and the asterisks next to species names flag taxonomic issues (see Appendix S1: Table S2). A unique element of the panel on the left is the sample size in parentheses, e.g. "(14,10)" for *Vanessa annabella*, which is the number of locations from which data were included from the NABA and Shapiro datasets, respectively. Finally, on the far right of panel (b), the lambda symbols represent the results of individual time series models run for the species present in the NABA program but without enough sites and years to be included in the main model (and thus not a part of the A list); a blue symbol indicates a species with an 80% or greater probability of increasing in recent years, while a red symbol indicates an 80% chance of decreasing, and black is neither increasing nor decreasing. The other species (beyond the top 50 highest ranked shown here) are included in Appendix S1: Figures S7, S8, and S9.

Without observational data, the ranking of B group species required a partitioning of weights among the other lines of information. To partly inform that process, we used the A group species to estimate the effects of other variables on risk index (based on NABA and Shapiro data). The model explained a relatively small proportion of variance in the risk index (Appendix S1: Table S1), but did demonstrate that smaller wingspans (99% probability of effect) and lower range-wide precipitation (98% probability of effect) are associated with risk for the A group species. In addition, we also suspected climate change would be important based on our previous work with western butterflies (Forister et al. 2021, Halsch et al. 2021). This is especially true given the large presence of B group species with ranges in the desert southwest (Appendix S1: Figure S4h), a region heavily impacted by warming and drying trends. We adopted the following weighting scheme to calculate a single risk value for species in the B group: 20% precipitation, 20% wingspan, 20% climate change, 10% development, 10% range size, 10% voltinism, and 10% host range; correlations among the seven variables as well as the two observational variables (for the A group) are shown in Appendix S1: Figure S6. As a comparison to that scheme, we also ranked the B group species with equal weights (14.3%) among the seven variables; the resulting risk values were correlated at *r* = 0.90 (*t* = 29.32, df = 210, *P* < 0.001) with the values from the primary scheme. With a third weighting scheme based on 50% from each of average range-wide precipitation and wingspan (the two variables identified as most predictive of risk by analysis of the A group), the correlation with the main scheme was *r* = 0.57 (*t* = 10.16, df = 210, *P* < 0.001).

The top fifty species with the highest risk values from each of the A and B groups are shown in Figure 2 (the other species with lower risk values are in Appendix S1: Figures S7, S8 and S9). For the highest-ranked A group species, agreement between the two monitoring schemes is apparent with large "risk circles" in both the NABA and Shapiro columns (Figure 3a). Time series plots for two of those top species are shown in Figure 3 (*Vanessa annabella*) and Figure 4 (*Euchloe ausonides*); in Figure 5, neutral or upward trajectories can be seen for *Poanes melane*, the species with the lowest risk index among the A group species (Appendix S1: Figure S10). Similar plots for all other A group species are available through an online tool (see Appendix S1). The rankings for the A group species are shown with 85% credible intervals (Figure 2a), which are broad; this uncertainty reflects the high inter-annual variability inherent to the time series data being modeled (from both NABA and Shapiro) and should be kept in mind when interpreting the position of species on the A group list.

**FIGURE 3.**
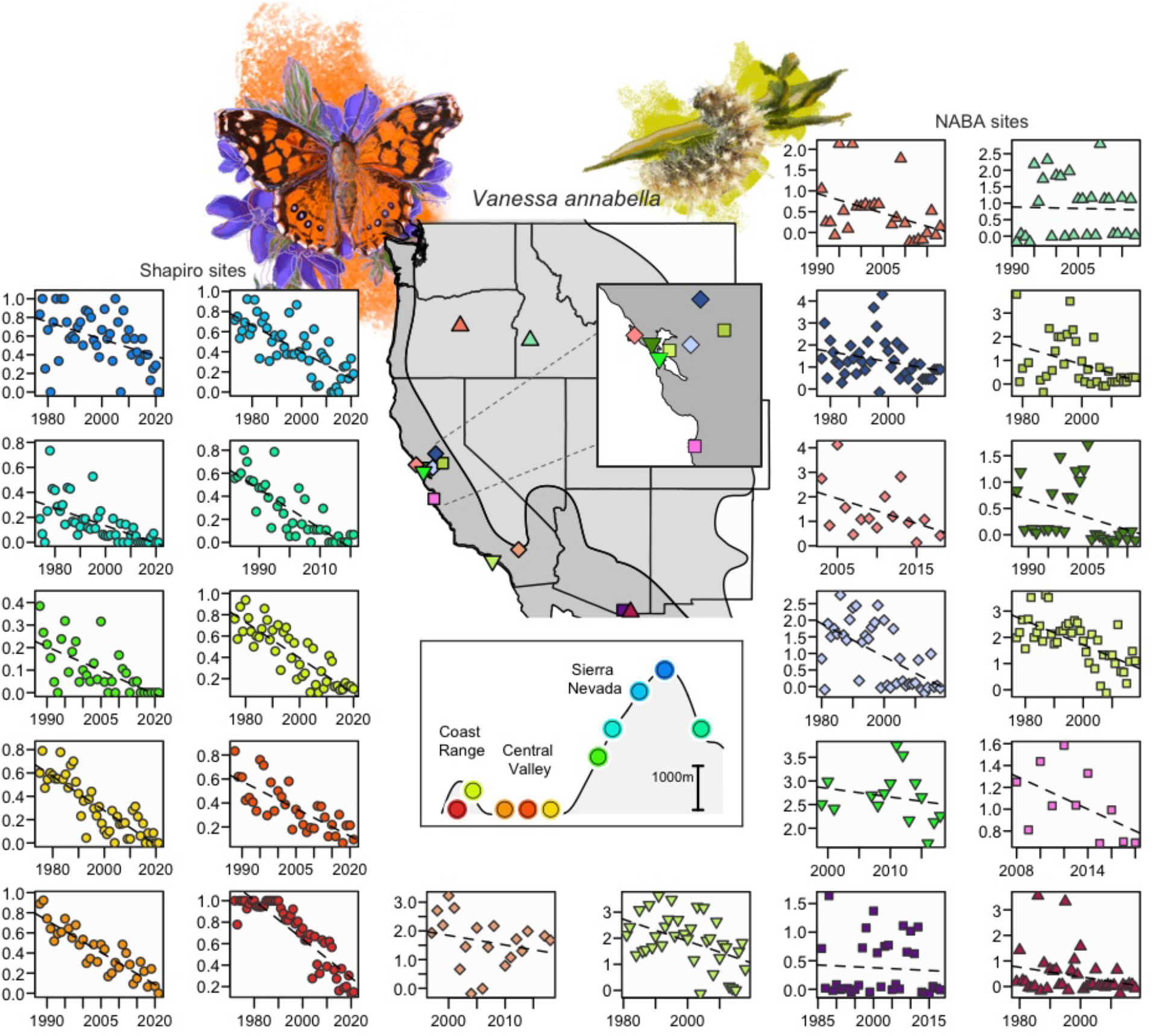
Overview of site-specific trends through time for *Vanessa annabella* at Shapiro sites (on the left) and NABA sites (on the right and along the bottom). Plots for Shapiro sites are shown with decreasing elevation and colored to match the elevational profile of Northern California (cooler colors are montane sites) shown below the map of the western US. The y-axes for Shapiro plots are the fraction of days a species was seen at a site in a year (Shapiro data were truncated at 1984 for analyses, but earlier years are shown here and in Figures 4 and 5). Plots for NABA sites are shown with decreasing latitude (starting with the most northern sites), from the top right to the bottom, with symbols matching the locations shown in the central map. Values shown in NABA plots have been adjusted for variation in sampling effort, and values plotted are total counts of individuals on a natural log scale. Also shown on the central map is the geographic range of *V. annabella* (Glassberg 2017), with the multivoltine portion of the range (closer to the coast) shown as darker gray. Adult and caterpillar images by Camryn Maher, copyright 2022.

**FIGURE 4.**
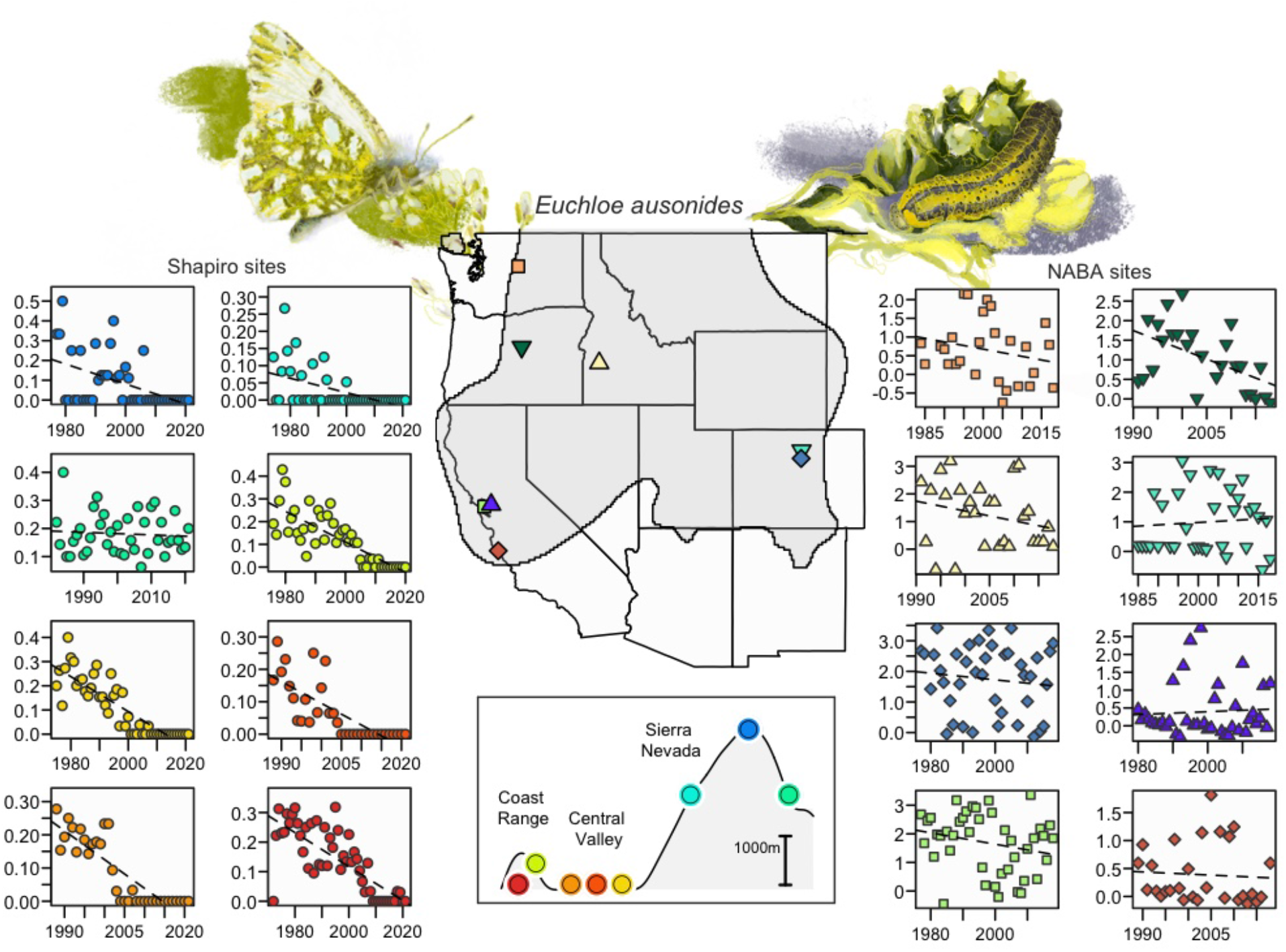
Overview of site-specific trends through time for *Euchloe ausonides* at Shapiro sites (on the left) and NABA sites (on the right). Plots for Shapiro sites are shown with decreasing elevation and colored to match the elevational profile of Northern California (cooler colors are montane sites) shown below the map of the western US. The y-axes for Shapiro plots are the fraction of days a species was seen at a site in a year. Plots for NABA sites are shown with decreasing latitude (starting with the most northern sites), with symbols matching the locations shown in the central map. Values shown in NABA plots have been adjusted for variation in sampling effort, and values plotted are total counts of individuals on a natural log scale. The geographic range of *E. ausonides* (Glassberg 2017) is shown as the gray shaded area on the central map. Adult and caterpillar images by Camryn Maher, copyright 2022.

**FIGURE 5.**
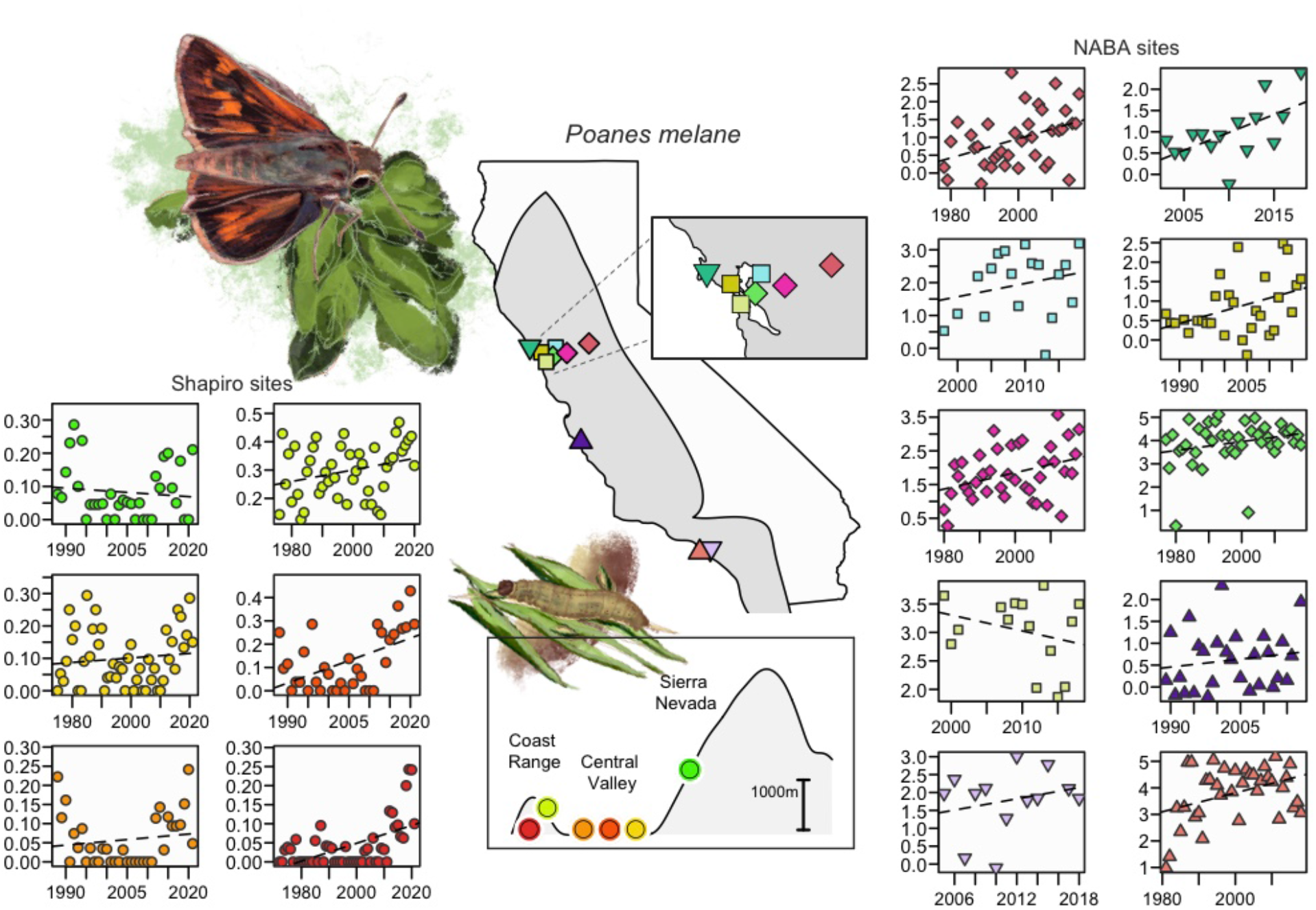
Overview of site-specific trends through time for *Poanes melane* at Shapiro sites (on the left) and NABA sites (on the right). Plots for Shapiro sites are shown with decreasing elevation (cooler colors are montane sites) and colored to match the elevational profile of Northern California shown below the map of the western US. The y-axes for Shapiro plots are the fraction of days a species was seen at a site in a year. Plots for NABA sites are shown with decreasing latitude (starting with the most northern sites), with symbols matching the locations shown in the central map and insect. Values shown in NABA plots have been adjusted for variation in sampling effort, and values plotted are total counts of individuals on a natural log scale. The geographic range of *P. melane* is shown as the gray shaded are on the central map. Adult and caterpillar images by Camryn Maher, copyright 2022.

It should also be remembered that the NABA data are based on a once-per-year census scheme, which potentially raises concerns with respect to phenological shifts: if species emerge earlier in response to warming springs, it is possible that a mid-summer count could erroneously infer decline if occurring past the population peak in more recent years. We took advantage of the temporally-intensive Shapiro program, where phenological shifts have been observed (Forister and Shapiro 2003), to simulate once-per-year sampling dates and compare results to a census based on biweekly counts. Contrary to the expectation that earlier emergence might lead to a mistaken inference of decline, but consistent with recent findings (Forister et al. 2018, Macgregor et al. 2019), earlier emergence tends to be associated with stable or increasing populations, and univoltine and bivoltine species do not appear as outliers in that relationship (Figure 6). Moreover, inferences about changes in population density over time are similar in once-per-year sampling and the full data with many visits per year (right column of panels in Figure 6). The role of phenological change in population response to anthropogenic stressors, including climate change, remains an important issue (Bonoan et al. 2021), but does not appear to bias the results presented here (see also Appendix S1: Figure S1 for a statistical control on phenology in the core NABA population model).

**FIGURE 6.**
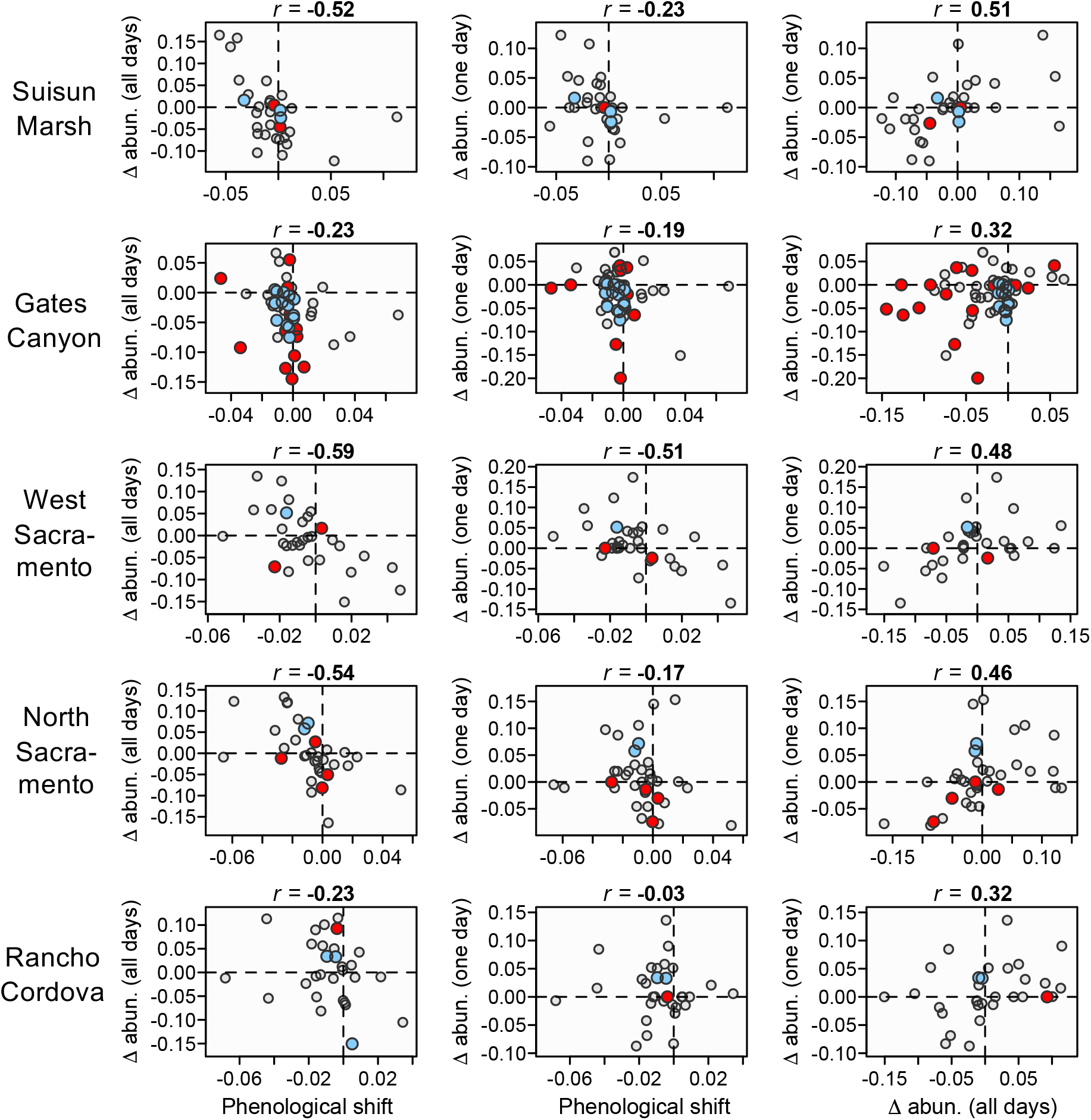
Analyses of phenology and abundance using Shapiro data and three variables: (1) change in annual abundance as summed across all visits within a year, shown as "Δ abun. (all days)"; (2) change in abundance based on a single, randomly sampled day each year, shown as "Δ abun. (one day)"; and (3) change in the date of first flight, shown as "Phenological shift". The values plotted are year coefficients for each species from models including year predicting each of the three variables, and Pearson correlation coefficients for each pairwise association are shown above the plots. Negative values for phenological shift indicate earlier emergence over time. For example, in the upper row, the negative relationship in the left panel indicates that species that emerged earlier (negative values on the x-axis) tended to become more abundant over time (positive values on the y-axis); and the same pattern is evident when change in abundance is estimated with the once-per-year sampling (middle panel); finally, in the right panel, it can be seen that change through time in abundance estimated with all of the data is positively correlated with year coefficients for change in abundance estimated with the once-per-year (NABA-style) samples. In all plots, red points are univoltine species with one generation per year, blue points are bivoltine species, and gray points represent species with more than two generations per year. Two outlier points (one species at Gates Canyon and one at Rancho Cordova) were strongly negative and compressed the visualize of other species; they are excluded here although the patterns and direction of relationships are unaltered.

Finally, we examined the distribution of the species-level risk index geographically and phylogenetically, for which we divided the species into low-, medium- and high-risk categories, based on composite risk values below the 75th quantile, between the 75th and 90th quantiles, and above the 90th, respectively. For the A group species, average forecast 50-year occupancy (based on NABA data) was 33.7% of populations extant (standard deviation = 12.5%) for the high-risk species, 47.7% occupancy (sd = 11.0%) for the medium-risk species, and 66.0% (sd = 13.2%) for the low-risk species. Considering the species with medium- and high-risk index values (above the 75th quantile of risk values) for the A group, the spread of average risk across the 11 western states is only partially associated with expected numbers of the most at-risk species (Figure 7a and 6c). For example, average risk is high in the northern Central Valley of California and in the northwestern region of Oregon (Figure 7a), while the cumulative occurrence of at-risk species is lower in those areas (Figure 7b). Similarly, the number of at-risk species (cumulative occurrence) is high in the Sierra Nevada, but average risk is lower. The distributions of risk for the B group species highlight the bias of that group towards the most southern areas, with high average risk along the southern California coast (Figure 7c) and a concentration of at-risk species along the border between Mexico and New Mexico (Figure 7d).

**FIGURE 7.**
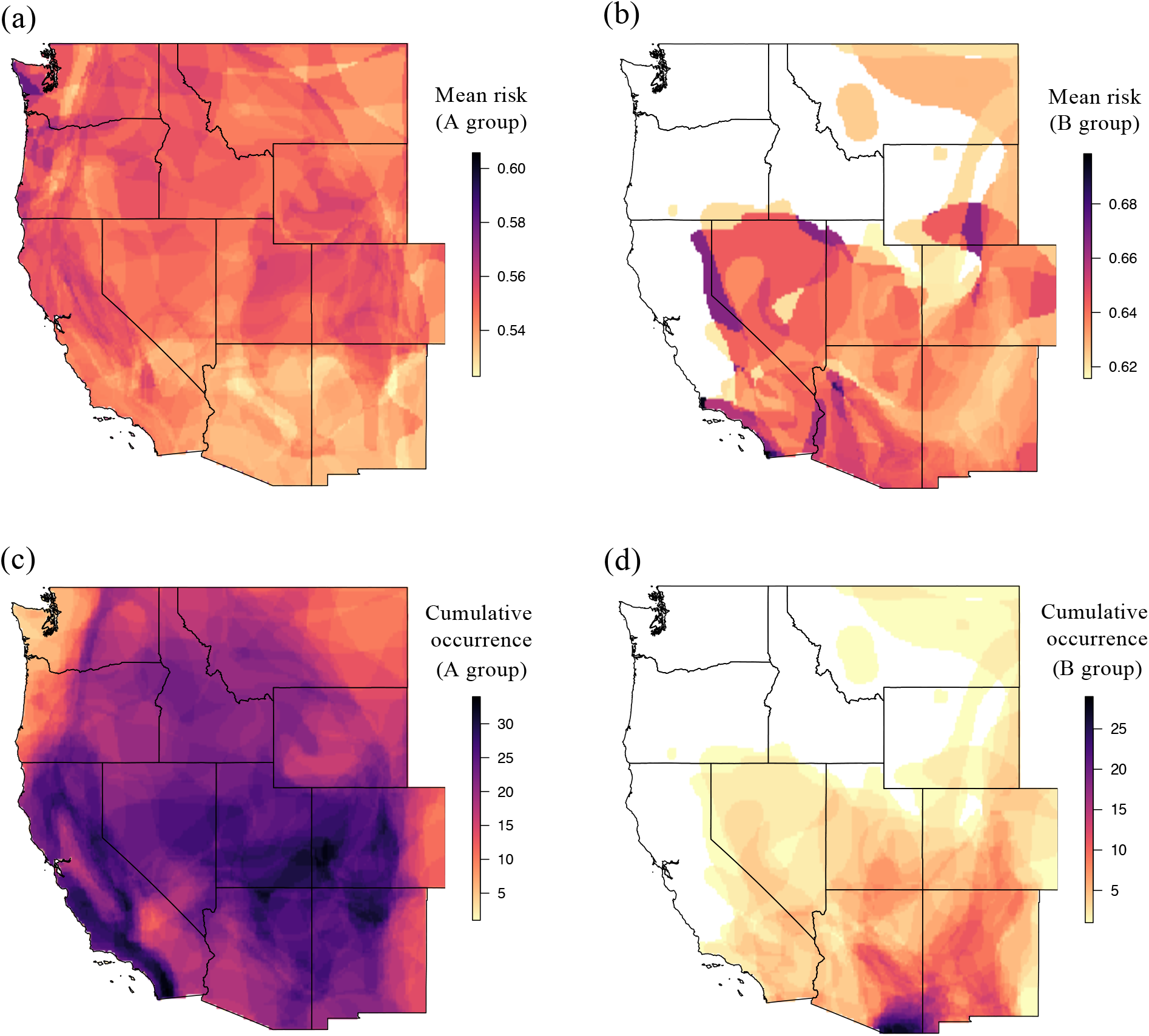
The geography of risk for species with values in the upper 75th quantile of risk indices as shown in Figure 2 (i.e., combining "medium" and "high" risk categories treated separately in Figure 8). Panels (a) and (b) show average risk values among those high risk species, separately for the A and B group species, while panels (c) and (d) show cumulative estimated species occurrence, again for the A group and B group species separately.

The phylogenetic picture of risk shows multiple clusters of at-risk species, and some lineages with notably lower risk, like much of the Nymphalidae (Figure 8). The families sharing the disproportionate amount of risk are the Lycaenidae (with 17% of species in the high-risk category, above the 90th quantile of risk) and the Hesperiidae (with 14% of species at high-risk); these are followed by the Riodinidae (with 13% of species at high risk, albeit based on a small sample size from a family represented by only 8 species) and the Pieridae (with 12% of species at high risk). The percentages of high-risk species in the Papilionidae and Nymphalidae are just 8% and 2%, respectively (Figure 8). Tests of phylogenetic inertia are consistent with the observation of phylogenetically clustered risk (Pagel’s λ = 0.38, *P* < 0.001; Blomberg’s *K* = 0.049, *P* = 0.001 based on 1k randomizations), which is evident not only at the family level, but also at lower taxonomic levels, including (among others) within the genera *Lycaena*, *Agathymus*, and *Eurema* (Figure 8).

**FIGURE 8.**
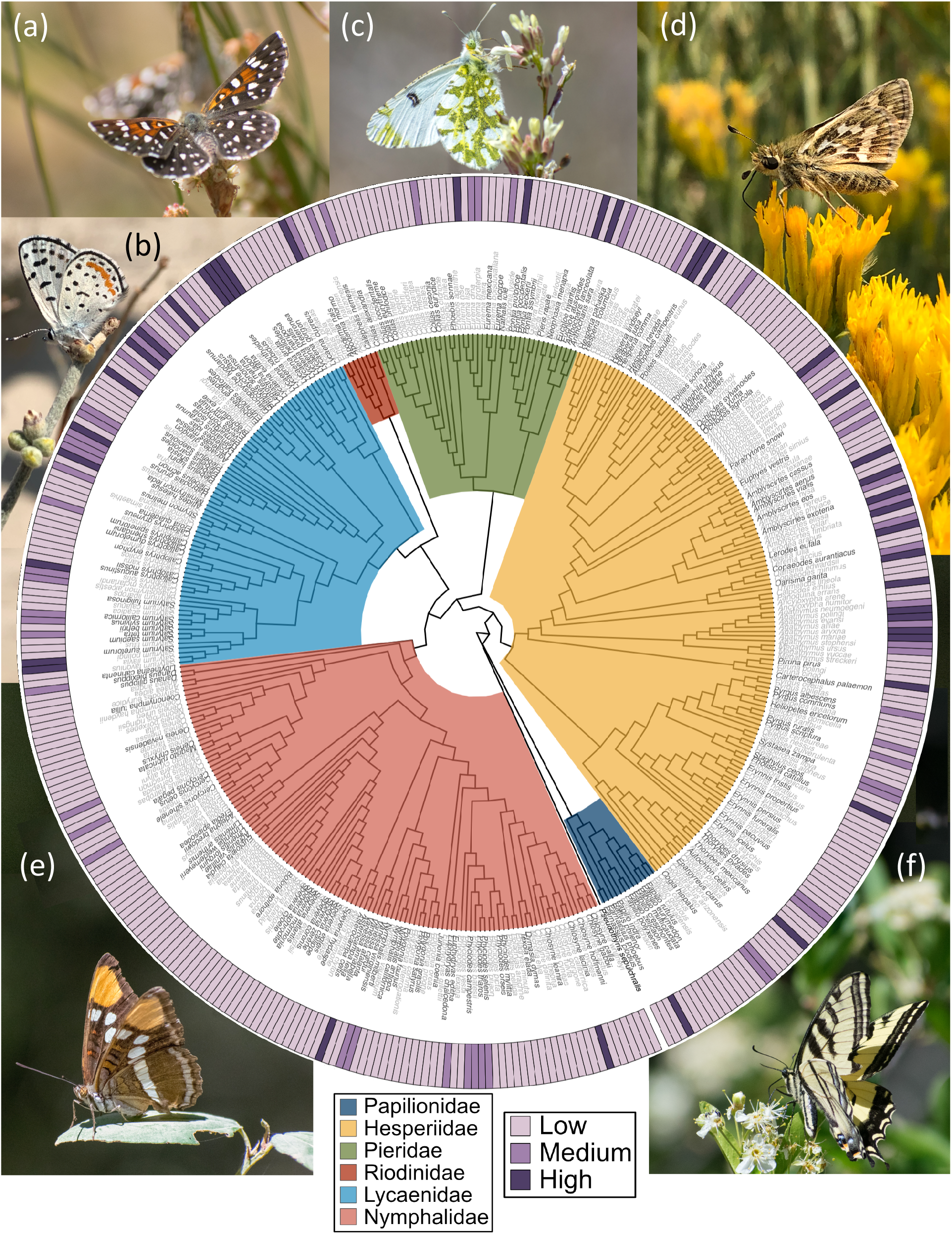
The phylogenetic distribution of risk, here shown as three categories: high risk (upper 90th quantile), medium risk (75th to 90th quantiles), and low risk (below the 75th quantile). Species names in black are the A group species, other are B group. Butterfly images as follows: (A) *Apodemia mormo* (Riodinidae); (B) *Euphilotes pallescens arenamontana* (Lycaenidae); (C) *Euchloe ausonides* (Pieridae); (D) *Polites sabuleti* (Hesperiidae); (E) *Adelpha bredowii* (Nymphalidae); (F) *Papilio rutulus* (Papilionidae). Photo credits go to CAH (panels A, C, E, and F); MLF (panels B and D). Bootstrap support is not shown but the vast majority of nodes have support above 0.95; see Zhang et al. (2019) for additional details.

## DISCUSSION

Our goal has been to organize and analyze heterogenous data sources in a way that allows conservation biologists to identify the butterflies in the 11 western US states that are most likely to suffer serious reductions in range or population size in coming years. It has not been our objective in this paper to document butterfly declines or to identify traits that make insects more or less sensitive to population stressors, as these topics have been addressed elsewhere for North America (Schultz et al. 2019, Wepprich et al. 2019, Forister et al. 2021), the Neotropics (Janzen and Hallwachs 2019, Salcido et al. 2020), and numerous other parts of the world (Nakamura 2011, Fox 2013, Wagner 2019). We hope that our work on species prioritization advances the issue for conservation practitioners using mixed data types with uneven spatial coverage and uncertainty in historical trends. Although some parts of the world (notably countries in Europe) have dense coverage with standardized monitoring, prioritization in most of the world will involve some mix of monitoring and trait-based inference.

The western states have been our region of study, rather than the entire US, because the impacts of climate change are severe and distinct in this arid region (Gonzalez et al. 2018), and the butterfly fauna is similarly shaped by a unique topography and climatic history (Shapiro 1996, Hawkins 2010). As a consequence of expansive areas with low human population density, about half of the butterfly species in the region are not included in the monitoring datasets used here, yet we have brought together information on the entire fauna (with the exception of a few species with rare occurrences, mostly strays across the US-Mexico border). Because of this, our study has an apples-and-oranges structure (species with and without monitoring data) that extends to the interpretation of the risk index values and engenders certain ironies. Chief among the ironies of our work is the fact that we rank B group species in part by certain variables (geographic range, exposure to climate change, etc.) that are not strongly associated with declines in the species for which we have historical records (the A group). In other words, considering Figure 2, the A group species near the top of the list do not necessarily have the smallest ranges, and the same can be said of other variables. Even for the two variables (wingspan and average precipitation) which do predict risk in the A group, the variance explained is low (Appendix S1: Table S1) yet we still emphasize these variables in ranking the B group species. We discuss these apparently counterintuitive decisions below, and then discuss phylogenetic and geographic hotspots of risk. Finally, we end with a consideration of individual taxa most deserving of attention given available evidence.

Among the complexities of variables potentially associated with risk, an understanding of geographic range starts by noting that the A group species have broader geographic ranges (Appendix S1: Figure S4a), which is part of the reason they are present at enough NABA sites to be included in our core population model. Thus the fact that many of the most severely declining species are widespread (e.g., *Vanessa annabella* in all 11 states) does not diminish the logic of prioritizing B group species based in part on small range size, which is a well-known determinant of risk (Staude et al. 2020). Similarly, the effects of voltinism and ecological host specialization are relatively straightforward: everything else being equal, we expect a species with multiple generations per year (a trait often associated with increased dispersal) and an ability to utilize many hosts to be more resilient to any number of stressors than another species without those traits (Eskildsen et al. 2015). We have previously observed the resiliency of multivoltine species during a mega-drought in the western US, where species with multiple generations per year were able to take advantage of earlier springs and greater time for population growth to temporarily reverse downward trajectories of multiple decades (Forister et al. 2018).

The interpretation of other variables is less straightforward, chief among them being exposure to climate change. Previous work with western butterflies has identified warming and drying conditions as stressors, based in particular on analyses of geographic variation among study sites in climate change effects and changes in aggregate butterfly density (Forister et al. 2021). At the species level (rather than the level of individual study sites), the same signal is not as apparent in the present study for the A group species (in other words, the species towards the top of the A group list do not have particularly high exposures to climate change). This is because most of these species have large enough ranges that their exposure to climate change (when quantified across the entire range) includes areas with both more and less severe warming and drying that tend to cancel each other out at the scale of broadly-distributed species. However, the B group species have smaller and more southern ranges (Appendix S1: Figure S4), which is the part of the west most impacted by climate change (Gonzalez et al. 2018). Thus, we believe exposure to climate change is well justified as a contributing factor to risk specifically for these species for which we lack monitoring data.

Exposure to development (urban, suburban and agricultural lands) requires similarly careful interpretation. This is chiefly because the data most well suited to understanding the effects of habitat destruction on insects will rarely be collected: places that have already been developed will not be monitored, and existing monitoring efforts will often be located in more pristine locations even when relatively proximate to human habitation (Wagner et al. 2021). The Shapiro dataset is an exception, as it encompasses a severe land use gradient from the agricultural and urban Central Valley to the undeveloped high elevations of the Sierra Nevada. From that program, we know that land conversion and contamination (with pesticides) have effects of similar magnitude at low elevations (Forister et al. 2016). Though similar information does not exist across the west, we included exposure to development in our rankings here for the B group species for the simple reason that common sense suggests that a range that encompasses more development is likely to experience increasing fragmentation and contamination in coming years relative to a species with less exposure.

Geographic projections of risk for B group species emphasize the southern areas of the west (Figure 7), but also point to specific hotpots of average risk that include the southern California coast. Like A group species in the Central Valley of California, that coastal region has a low cumulative occurrence of B group species, but on average the species that are there in the vicinity of the Los Angeles basin score high for our risk factors. Arizona and southwestern New Mexico have a high concentration of B group species with high risk factors, thus this area should be prioritized for future monitoring efforts. For A group species, the Sierra Nevada Mountains (especially the northern Sierra), the Colorado Plateau and the southern Rocky Mountains are hotspots of declining species (Figure 7). These same places have been recently identified as hotspots of imperiled species in analyses that included plants, vertebrates, freshwater invertebrates and some terrestrial insects (Hamilton et al. 2022).

Phylogenetically, risk values are strongly clustered within and among families, with notable concentrations in the Lycaenidae and Hesperiidae, with the latter in part due to both species with small southern ranges (B group species) and species in monitoring programs with observed declines. The phylogenetic clustering of risk suggests that currently unknown or unmeasured variables could in the future improve our ability to model interspecific variation in populations trajectories and extinction risk. On the other hand, the non-random distribution of risk among species and lineages suggests that species loss might itself be clustered, leading to shifts in the function and composition of assemblages (Sol et al. 2017). At present, we can say that of the high-risk category species (with risk index values above the 90th quantile), 46% are Hesperiidae. The family Nymphalidae has the lowest concentration of at-risk species, although one of the most notably declining species is in this family. Despite being large and dispersive and able to use a number of exotic plants as larval hosts, *Vanessa annabella* is becoming hard to find across locations that include urban centers, high mountains, and southern deserts (Figure 3).

Although *V. annabella* is deservedly at the top of the risk list (Figures 2 and 3), we stress the uncertainty in the actual risk values that we have generated, and we do not place much weight on the exact position of species on that list. In other words, we believe that the top species in the A group are indeed in historical declines that will likely continue in coming years, but the fact that one species is in the 4th position vs the 10th or even the 25th position on the list is not necessarily important. Small differences in, for example, the projected 50-year probability of population persistence affect the positions for those top species which have mostly similar risk values (and broadly overlapping credible intervals). This is why we conservatively suggest that all of the top 50 species in the A group (Figure 2) deserve closer scrutiny and in some cases likely deserve protection. The fact that rankings should be treated as approximate is also why we have presented other lines of information (geographic range, host specialization, etc.) for the A group, even though the risk index ranking is based solely on the observational data (NABA and Shapiro) for those species. For example, *Pontia protodice* and *Lycaena xanthoides* have nearly identical risk indices, but the latter (*L. xanthoides*) is univoltine with a smaller geographic range, greater exposure to development and a more specialized diet (Figure 2); these are all factors that could be considered by conservation biologists and ecologists interested in declining insects. With respect to current protections, only two of the species that we have studied have status at the federal level: one of the A group species (the monarch butterfly, *Danaus plexippus*) is currently a candidate for protection under the US Endangered Species Act (ESA), and one of the B group species, *Lycaena hermes*, is currently listed as threatened.

Our presentation of the top 50 species in the A group (Figure 2) includes sample sizes (for NABA and Shapiro datasets) which should also be considered when judging the evidence for risk. For example, the 2nd and 3rd species on the A group list (Figure 2) are represented by data from 3 or fewer sites for the NABA and Shapiro datasets. The small samples for those species are reflected in broad intervals around the risk values, and it can be noted that other species in the top 10 for the A list are known to be in decline based on evidence from two to three times as many sites (e.g., *Pholisora catullus*, *Atalopedes campestris*, and *Euchloe ausonides*). The number of sites for individual species is a reflection not just of information available for analysis. It should be remembered that risk associated with the NABA data derives from a multi-species population viability analysis. In that analysis, species with fewer sites are more likely by chance to have lower occupancy in forecasts than species known from a greater number of sites. This is both a methodological feature of stochastic simulations but also reflects a biological reality in that more widespread species are known from a greater number of NABA sites (thus geographic range is indirectly involved in the contribution that the NABA analyses make to our estimate of risk).

Yet another important aspect of sample size involves A group species not represented in both of the observational datasets; for these species, we used median interpolation. In other words, when calculating the risk index for a species present in, for example, the Shapiro dataset but not NABA, we assigned a 50-year projection value based on the median across all other species represented in the NABA dataset. For the present effort, we consider this to be at least a relatively simple assumption, although we acknowledge that future analyses could use more sophisticated interpolation perhaps including information from closely related species. The phylogenetic signal observed here suggests that genetic relatedness could be a tool for dealing with uncertainty and missing data in conservation ranking.

The weight of missing data and uncertainty of course becomes greater when we turn to the top 50 species in the B group (Figure 2) for which monitoring data is either absent or insufficient for robust models. Not only is robust observational data lacking, but so many of the B group species are similar in having small ranges in hot and dry parts of the region that the overall spread of risk values is smaller than for the A group. Thus, rankings in the top 50 for the B group should be treated as approximate. For example, *Strymon avalona* is restricted entirely to Catalina Island (less than 200 square kilometers) off the coast of southern California. The partly wild nature of the island gives the species a low development score and the area happens to be characterized by only moderate departure from climatic baseline. Thus *S. avalona* ranks towards the bottom of the top 50 for the B group (Figure 2), even though that small geographic range of course puts it at high risk of stochastic loss. Similarly, many of the B group species below the top 50 have negative annual trends (indicated by red lambda symbols to the right of the panel in Appendix S1: Figures S7 - S9), albeit based on very few NABA sites (which is why we have shown those results but did not use them in the calculation of the B group risk index). In general we hope that the data organized here for the B group species is an inspiration for greater monitoring of these taxa with small ranges in regions vulnerable to threats that include ongoing climate change and the loss of natural disturbance regimes (Haddad 2018).

## CAVEATS & CONCLUSIONS

Our synthesis of status and trends for a diverse fauna faced many challenges, and includes many sources of taxonomic and spatial bias in the data that were available to us. We have not undertaken a formal assessment of bias for the temporal and other patterns reported here (Boyd et al. 2022), largely because the number of datasets to be assessed is large and the issue deserves another manuscript-level treatment. In addition to sources of spatial bias discussed above, including the over-representation of widespread species in monitoring programs, we close by noting that even for those widespread species (well represented in census data), the information tends to be clustered around areas of human population density. Thus, broad ranges (e.g., Figure 3) and relatively more narrow ranges (e.g., Figure 5) alike are not particularly well sampled in terms of the spread of monitored locations in space. We hope that these results inspire greater investment in state-level monitoring programs (Taron and Ries 2015, Wepprich et al. 2019), which could eventually fill data gaps and lead to a national understanding of butterfly status on par with countries in Europe. Coming years should also see the development of new models that can take advantage of mixed data types including those reported from crowd-sourced platforms (e.g, Strebel et al. 2022). Our estimation of risk has not included exposure to pesticides, which is available at the county level for California, but not for other states in the region, although we know that it is an important stressor (Gilburn et al. 2015, Forister et al. 2016).

Another important issue that we acknowledge is that our estimates of exposure to development and climate change are restricted to the portions of geographic ranges found in the 11 western states. This was mainly motivated by our focus on the unique exposure of the region to warming and drying trends, but it is of course the case that wide ranging species might have other parts of their range subject to divergent pressures. Future analyses of risk could quantify heterogeneity of stressors at continental scales. In the meantime, it is for these reasons that we have included our qualitative range labels (N, S, E, and W) with our rankings (Figure 2), which the reader can use to focus as desired on species as a function of their distribution.

The traditional focus for butterfly conservation in the United States has been at the taxonomic level of subspecies, which is partly a consequence of the fact that population segments cannot be listed for invertebrates (thus leaving subspecies as the next unit below full species that can be protected). Thus, we acknowledge that our results fall partly outside of the traditional scope of conservation work for butterflies in the United States. It is, however, entirely likely that compounding population losses across the wild spaces of the region have pushed many full species to the point where range-wide research and conservation attention are warranted. A notable example of this is recent effort focused on conservation of the monarch butterfly, *Danaus plexippus* (Pelton et al. 2019), which is indeed in our list of the 50 most at-risk species (Figure 3). Notably, a large number of species are higher on the list and are equally deserving of attention. Research in coming years might also profitably focus on species that appear to be relatively stable. One example (*Poanes melane*) is shown in Figure 5, and others can be found toward the end of the ranked list of species in Appendix S1: Figure S10, including *Ochlodes agricola, Papilio rutulus*, and *Limenitis lorquini*. We hesitate to use the common metaphor of winners and losers. That implies that the game is over, when of course the Anthropocene is underway. Nevertheless, the diversity of ecologies, morphologies and geographic ranges among the stable or increasing species (Appendix S1: Figure S10) suggests that much could be learned about combinations of traits potentially associated with resilience. It is our hope that the results presented here are a framework that will facilitate such work in coming decades while acknowledging the many assumptions that have been made along the way to synthesize diverse data and organize species by composite risk scores.

## ACKNOWLEDGEMENTS

MLF thanks the National Science Foundation (DEB-2114793), and CAH was supported by a National Institute of Food and Agriculture fellowship (NEVW-2021-09427). EMG, KLB, and JPJ were supported by the Modelscape Consortium with funding from NSF (OIA-2019528). Thanks to the Plant Insect Group at UNR for much thoughtful feedback, and Lee Dyer in particular for various key ideas, as well as Sarina Jepsen who provided important feedback on subspecies risk. We thank Texas State University for the use of the LEAP computing cluster, and thanks also to authors of our previous analysis of western butterflies (Forister et al. 2021) without which the current paper would not have gotten off the ground.

## CONFLICT OF INTEREST

The authors declare no conflict of interest.

## AUTHOR CONTRIBUTIONS

MLF conceived the project and the overall organization of data and the presentation of results. TVR wrote the Bayesian population models for the NABA data and contributed to the design of analyses. EMG contributed to numerous components, especially the analysis of spatial data. KJB organized information on subspecies. CAH, CFC, KLB, and TB contributed to component analyses and assisted with data collation or organization. JZ, QC, and NVG generated the phylogeny which was analyzed here by JPJ. JG manages the collection of NABA data and contributed the expert-derived range outlines. AMS has collected the vast majority of the Shapiro transect data (with a few recent years of collection contributed in part by MLF and CAH). All authors contributed to interpretation of results and writing of the manuscript.

## DATA AVAILABILITY STATEMENT

Data analyzed for this paper are partly based on data used in Forister et al. (2021), with additions and updates. The North American Butterfly Association (NABA) dataset used here is the same as in Forister et al. (2021), and is available at https://doi.org/10.5281/zenodo.4460647. The Shapiro transect dataset has been updated and will be archived on figshare.com upon acceptance for publication (during review the dataset is available at https://sites.google.com/view/westernbutterflies/data). New variables used here include: exposure to land development, which derives from the 2020 Cropland Data Layer (USDA 2020) downloaded using code from the function getCDL from the cdlTools R package (Chen 2018), and climate data, including exposure to climate change, which derive from TerraClimate (Abatzoglou et al. 2018) downloaded from THREDDS (Unidata 2021a) using NCSS (Unidata 2021b) with batch scripts following the example developed by Katherine Hegewisch (https://climate.northwestknowledge.net/TERRACLIMATE/pages/guidance/ CODE/terraclimate_ncss_subset.sh). Those variables for climate and land use will be included upon manuscript acceptance in the same figshare.com archive as the Shapiro dataset, along with the following variables that are updated, relative to data used in Forister et al. (2021): geographic ranges, voltinism, diet breadth, and wingspan. Geographic ranges and voltinism derive from Glassberg (2017, 2018) and from Scott (1986). Diet breadth data were taken primarily from Scott (1986) but also from Brock and Kaufman (2006) and Lotts et al. (2007); wingspan data derive from Opler (1999), supplemented with Warren et al. (Warren et al. 2013). Code used is not novel to this project (relevant functions and packages are cited).

## Notes

### Competing Interest Statement

The authors have declared no competing interest.

### Summary of Updates

In this revision, part of the supplemental material has been moved into the main text (mainly figure 6) in response to reviewer suggestions.

## REFERENCES

Abatzoglou, J. T., S. Z. Dobrowski, and S. A. Parks. 2020. Multivariate climate departures have outpaced univariate changes across global lands. Scientific Reports 10:1–9.

Abatzoglou, J. T., S. Z. Dobrowski, S. A. Parks, and K. C. Hegewisch. 2018. TerraClimate, a high-resolution global dataset of monthly climate and climatic water balance from 1958–2015. Scientific Data 5:170191.

Altschul, S. F., T. L. Madden, A. A. Schäffer, J. Zhang, Z. Zhang, W. Miller, and D. J. Lipman. 1997. Gapped BLAST and PSI-BLAST: a new generation of protein database search programs. Nucleic acids research 25:3389–3402.

Benton, T. G. 2003. Understanding the ecology of extinction: are we close to the critical threshold? Annales Zoologici Fennici 40:71–80.

Bladon, A., R. Smith, and W. Sutherland. 2022. Butterfly and moth conservation: global evidence for the effects of interventions for butterflies and moths. University of Cambridge, Cambridge, UK.

Bonelli, S., L. P. Casacci, F. Barbero, C. Cerrato, L. Dapporto, V. Sbordoni, S. Scalercio, A. Zilli, A. Battistoni, and C. Teofili. 2018. The first red list of Italian butterflies. Insect Conservation and Diversity 11:506–521.

Bonoan, R. E., E. E. Crone, C. B. Edwards, and C. B. Schultz. 2021. Changes in phenology and abundance of an at-risk butterfly. Journal of Insect Conservation 25:499–510.

Boyd, R. J., G. D. Powney, F. Burns, A. Danet, F. Duchenne, M. J. Grainger, S. G. Jarvis, G. Martin, E. B. Nilsen, and E. Porcher. 2022. ROBITT: A tool for assessing the risk-of-bias in studies of temporal trends in ecology. Methods in Ecology and Evolution 13:1497–1507.

Brock, J. P., and K. Kaufman. 2006. Kaufman field guide to butterflies of North America. Houghton Mifflin Harcourt.

Brook, B. W., N. S. Sodhi, and C. J. A. Bradshaw. 2008. Synergies among extinction drivers under global change. Trends in Ecology & Evolution 23:453–460.

Cardoso, P., P. S. Barton, K. Birkhofer, F. Chichorro, C. Deacon, T. Fartmann, C. S. Fukushima, R. Gaigher, J. C. Habel, and C. A. Hallmann. 2020. Scientists’ warning to humanity on insect extinctions. Biological Conservation 242:108426.

Cardoso, P., T. L. Erwin, P. A. V Borges, and T. R. New. 2011. The seven impediments in invertebrate conservation and how to overcome them. Biological Conservation 144:2647– 2655.

Casner, K., M. L. Forister, K. Ram, and A. M. Shapiro. 2014a. The utility of repeated presence-absence data as a surrogate for counts: a case study using butterflies. Ecological Applications 18:13–27.

Casner, K. L., M. L. Forister, J. M. O’Brien, J. H. Thorne, D. P. Waetjen, and A. M. Shapiro. 2014b. Loss of agricultural land and a changing climate contribute to decline of an urban butterfly fauna. Conservation Biology 28:773–782.

Chen, L. 2018. cdlTools: Tools to Download and Work with USDA Cropscape Data. R package version 0.13.

Collins, T. J. 2007. ImageJ for microscopy. Biotechniques 43:S25–S30.

Diniz-Filho, J. A. F., P. De Marco Jr, and B. A. Hawkins. 2010. Defying the curse of ignorance: perspectives in insect macroecology and conservation biogeography. Insect Conservation and Diversity 3:172–179.

Dirzo, R., H. S. Young, M. Galetti, G. Ceballos, N. J. B. Isaac, and B. Collen. 2014. Defaunation in the Anthropocene. Science 345:401–406.

Edge, D. A., and S. Mecenero. 2015. Butterfly conservation in southern Africa. Journal of Insect Conservation 19:325–339.

Eisenhauer, N., A. Bonn, and C. A Guerra. 2019. Recognizing the quiet extinction of invertebrates. Nature Communications 10:1–3.

Eskildsen, A., L. G. Carvalheiro, W. D. Kissling, J. C. Biesmeijer, O. Schweiger, and T. T. Høye. 2015. Ecological specialization matters: long-term trends in butterfly species richness and assemblage composition depend on multiple functional traits. Diversity and Distributions 21:792–802.

Farber, O., and R. Kadmon. 2003. Assessment of alternative approaches for bioclimatic modeling with special emphasis on the Mahalanobis distance. Ecological Modelling 160:115–130.

Forister, M. L., B. Cousens, J. G. Harrison, K. Anderson, J. H. Thorne, D. Waetjen, C. C. Nice, M. De Parsia, M. L. Hladik, and R. Meese. 2016. Increasing neonicotinoid use and the declining butterfly fauna of lowland California. Biology Letters 12:20160475.

Forister, M. L., J. A. Fordyce, C. C. Nice, J. H. Thorne, D. P. Waetjen, and A. M. Shapiro. 2018. Impacts of a millennium drought on butterfly faunal dynamics. Climate Change Responses 5:3.

Forister, M. L., C. A. Halsch, C. C. Nice, J. A. Fordyce, T. E. Dilts, J. C. Oliver, K. L. Prudic, A. M. Shapiro, J. K. Wilson, and J. Glassberg. 2021. Fewer butterflies seen by community scientists across the warming and drying landscapes of the American West. Science 371:1042–1045.

Forister, M. L., A. C. McCall, N. J. Sanders, J. A. Fordyce, J. H. Thorne, J. O’Brien, D. P. Waetjen, and A. M. Shapiro. 2010. Compounded effects of climate change and habitat alteration shift patterns of butterfly diversity. Proceedings of the National Academy of Sciences of the United States of America 107:2088–2092.

Forister, M. L., and A. M. Shapiro. 2003. Climatic trends and advancing spring flight of butterflies in lowland California. Global Change Biology 9:1130–1135.

Fox, R. 2013. The decline of moths in Great Britain: a review of possible causes. Insect Conservation and Diversity 6:5–19.

Fox, R., M. S. Warren, T. M. Brereton, D. B. Roy, and A. Robinson. 2011. A new Red List of British butterflies. Insect Conservation and Diversity 4:159–172.

Franke, S., S. Pinkert, R. Brandl, and S. Thorn. 2022. Modeling the extinction risk of European butterflies and odonates. Ecology and Evolution 12:e9465.

Geyle, H. M., M. F. Braby, M. Andren, E. P. Beaver, P. Bell, C. Byrne, M. Castles, F. Douglas, R. V Glatz, and B. Haywood. 2021. Butterflies on the brink: identifying the Australian butterflies (Lepidoptera) most at risk of extinction. Austral Entomology 60:98–110.

Gilburn, A. S., N. Bunnefeld, J. M. Wilson, M. S. Botham, T. M. Brereton, R. Fox, and D. Goulson. 2015. Are neonicotinoid insecticides driving declines of widespread butterflies? PeerJ 3:e1402.

Glassberg, J. 2017. A Swift guide to butterflies of North America. 2nd edition. Princeton University Press.

Glassberg, J. 2018. A Swift guide to butterflies of Mexico and Central America. 2nd edition. Princeton University Press.

Gonzalez, P., F. Wang, M. Notaro, D. J. Vimont, and J. W. Williams. 2018. Disproportionate magnitude of climate change in United States national parks. Environmental Research Letters 13:104001.

Goulson, D. 2019. The insect apocalypse, and why it matters. Current Biology 29:R967–R971.

Haddad, N. M. 2018. Resurrection and resilience of the rarest butterflies. PLoS Biology 16:e2003488.

Hallmann, C. A., M. Sorg, E. Jongejans, H. Siepel, N. Hofland, H. Schwan, W. Stenmans, A. Müller, H. Sumser, T. Hörren, and others. 2017. More than 75 percent decline over 27 years in total flying insect biomass in protected areas. PlOS ONE 12:e0185809.

Halsch, C. A., A. M. Shapiro, J. A. Fordyce, C. C. Nice, J. H. Thorne, D. P. Waetjen, and M. L. Forister. 2021. Insects and recent climate change. Proceedings of the National Academy of Sciences:10.1073/pnas.2002543117.

Hamilton, H., R. L. Smyth, B. E. Young, T. G. Howard, C. Tracey, S. Breyer, D. R. Cameron, A. Chazal, A. K. Conley, and C. Frye. 2022. Increasing taxonomic diversity and spatial resolution clarifies opportunities for protecting imperiled species in the US. Ecological Applications: e2534.

Hawkins, B. A. 2010. Multiregional comparison of the ecological and phylogenetic structure of butterfly species richness gradients. Journal of Biogeography 37:647–656.

Hijmans, R. J., J. van Etten, M. Mattiuzzi, M. Sumner, J. A. Greenberg, O. P. Lamingueiro, A. Bevan, E. B. Racine, and A. Shortridge. 2021. raster: Geographic Data Analysis and Modeling, Version 2.9–23, R package.

Hughes, J. B., G. C. Daily, and P. R. Ehrlich. 2000. Conservation of insect diversity: a habitat approach. Conservation Biology 14:1788–1797.

Jamwal, P. S., M. Di Febbraro, M. L. Carranza, M. Savage, and A. Loy. 2021. Global change on the roof of the world: Vulnerability of Himalayan otter species to land use and climate alterations. Diversity and Distributions.

Janzen, D. H., and W. Hallwachs. 2019. Perspective: Where might be many tropical insects? Biological Conservation 233:102–108.

Katoh, K., K. Misawa, K. Kuma, and T. Miyata. 2002. MAFFT: a novel method for rapid multiple sequence alignment based on fast Fourier transform. Nucleic Acids Research 30:3059–3066.

Kellner, K. 2017. R Package ‘jagsUI’: a wrapper around ‘rjags’ to streamline ‘JAGS’Analyses, v.1.4.9.

van Klink, R., D. E. Bowler, K. B. Gongalsky, A. B. Swengel, A. Gentile, and J. M. Chase. 2020. Meta-analysis reveals declines in terrestrial but increases in freshwater insect abundances. Science 368:417–420.

Li, L., C. J. Stoeckert, and D. S. Roos. 2003. OrthoMCL: identification of ortholog groups for eukaryotic genomes. Genome Research 13:2178–2189.

Lotts, K. C., T. Naberhaus, and E. Sellers. 2007. PS 72–135: The butterflies and moths of North America: A database for research, education, and conservation.

Macgregor, C. J., C. D. Thomas, D. B. Roy, M. A. Beaumont, J. R. Bell, T. Brereton, J. R. Bridle, C. Dytham, R. Fox, and K. Gotthard. 2019. Climate-induced phenology shifts linked to range expansions in species with multiple reproductive cycles per year. Nature Communications 10:4455.

Maes, D., W. Vanreusel, I. Jacobs, K. Berwaerts, and H. Van Dyck. 2012. Applying IUCN Red List criteria at a small regional level: a test case with butterflies in Flanders (north Belgium). Biological Conservation 145:258–266.

McLaughlin, J. F., J. J. Hellmann, C. L. Boggs, and P. R. Ehrlich. 2002. Climate change hastens population extinctions. Proceedings of the National Academy of Sciences of the United States of America 99:6070–6074.

NABA. 2018. Checklist of North American Butterflies Occurring North of Mexico - Edition 2.4.

Naeem, S., R. Chazdon, J. E. Duffy, C. Prager, and B. Worm. 2016. Biodiversity and human well-being: an essential link for sustainable development. Proceedings of the Royal Society B: Biological Sciences 283:20162091.

Nakamura, Y. 2011. Conservation of butterflies in Japan: status, actions and strategy. Journal of Insect Conservation 15:5–22.

New, T. R., R. M. Pyle, J. A. Thomas, C. D. Thomas, and P. C. Hammond. 1995. Butterfly Conservation Management. Annual Review of Entomology 40:57–83.

Nguyen, L.-T., H. A. Schmidt, A. Von Haeseler, and B. Q. Minh. 2015. IQ-TREE: a fast and effective stochastic algorithm for estimating maximum-likelihood phylogenies. Molecular Biology and Evolution 32:268–274.

Nice, C. C., M. L. Forister, Z. Gompert, J. A. Fordyce, and A. M. Shapiro. 2014. A hierarchical perspective on the diversity of butterfly species’ responses to weather in the Sierra Nevada Mountains. Ecology 95:2155–2168.

Opler, P. A. 1999. A field guide to western butterflies. Houghton Mifflin Harcourt.

Pelham, J. P. 2022. A catalogue of the butterflies of the United States and Canada.

Pelton, E. M., C. B. Schultz, S. J. Jepsen, S. H. Black, and E. E. Crone. 2019. Western monarch population plummets: status, probable causes, and recommended conservation actions. Frontiers in Ecology and Evolution 7:258.

R Core Team. 2020. R: A language and environment for statistical computing R Foundation for Statistical Computing, Vienna, Austria. URL https://www.R-project.org/.

Revell, L. J. 2012. phytools: an R package for phylogenetic comparative biology (and other things). Methods in Ecology and Evolution:217–223.

Riecke, T. V, D. Gibson, M. Kéry, and M. Schaub. 2021. Sharing detection heterogeneity information among species in community models of occupancy and abundance can strengthen inference. Ecology and evolution 11:18125–18135.

Salcido, D. M., M. L. Forister, H. G. Lopez, and L. A. Dyer. 2020. Ecosystem services at risk from declining taxonomic and interaction diversity in a tropical forest. Scientific Reports 10:1–10.

Samways, M. J. 2007. Insect conservation: a synthetic management approach. Annu. Rev. Entomol. 52:465–487.

Schultz, C. B., N. M. Haddad, E. H. Henry, and E. E. Crone. 2019. Movement and demography of at-risk butterflies: building blocks for conservation. Annual Review of Entomology 64:167–184.

Scott, J. A. 1986. The butterflies of North America. Stanford University Press, Stanford, California.

Sekar, S. 2012. A meta-analysis of the traits affecting dispersal ability in butterflies: can wingspan be used as a proxy? Journal of Animal Ecology 81:174–184.

Sen, P. K. 1968. Estimates of the regression coefficient based on Kendall’s tau. Journal of the American Statistical Association 63:1379–1389.

Shapiro, A. M. 1996. Status of butterflies. Pages 743–757 Sierra Nevada Ecosystem Project: Final Report to Congress Vol. II. Center for Water and Wildland Resources, University of California, Davis, Davis, California.

Shapiro, A. M. 2022. Art Shapiro’s Butterfly Site https://butterfly.ucdavis.edu/.

Sol, D., I. Bartomeus, C. González-Lagos, and S. Pavoine. 2017. Urbanisation and the loss of phylogenetic diversity in birds. Ecology Letters 20:721–729.

Staude, I. R., L. M. Navarro, and H. M. Pereira. 2020. Range size predicts the risk of local extinction from habitat loss. Global Ecology and Biogeography 29:16–25.

Strebel, N., M. Kéry, J. Guélat, and T. Sattler. 2022. Spatiotemporal modelling of abundance from multiple data sources in an integrated spatial distribution model. Journal of Biogeography 49:563–575.

van Swaay, C., D. Maes, S. Collins, M. L. Munguira, M. Šašić, J. Settele, R. Verovnik, M. Warren, M. Wiemers, and I. Wynhoff. 2011. Applying IUCN criteria to invertebrates: How red is the Red List of European butterflies? Biological Conservation 144:470–478.

Taron, D., and L. Ries. 2015. Butterfly monitoring for conservation. Pages 35–57 Butterfly Conservation in North America. Springer.

Theil, H. 1950. A rank-invariant method of linear and polynomial regression analysis. Indagationes Mathematicae 12:173.

Turvey, S. T., and J. J. Crees. 2019. Extinction in the Anthropocene. Current Biology 29:R982– R986.

Unidata. 2021a. THREDDS Data Server (TDS) version 5.0. Boulder, CO: UCAR/Unidata.

Unidata. 2021b. NetCDF Subset Service. Boulder, CO: UCAR/Unidata.

USDA. 2020. National Agricultural Statistics Service Cropland Data Layer. Published crop-specific data layer. https://nassgeodata.gmu.edu/CropScape/.

Wagner, D. L. 2019. Insect declines in the Anthropocene. Annual Review of Entomology 65.

Wagner, D. L., E. M. Grames, M. L. Forister, M. R. Berenbaum, and D. Stopak. 2021. Insect decline in the Anthropocene: Death by a thousand cuts. Proceedings of the National Academy of Sciences 118:e2023989118.

Warren, A. D., K. J. Davis, E. M. Stangeland, J. P. Pelham, and N. V Grishin. 2013. Illustrated lists of American butterflies [21-XI-2017]. https://www.butterfliesofamerica.com/L/Neotropical.htm.

Wepprich, T., J. R. Adrion, L. Ries, J. Wiedmann, and N. M. Haddad. 2019. Butterfly abundance declines over 20 years of systematic monitoring in Ohio, USA. PLOS ONE 14:e0216270.

Wilson, E. O. 1987. The little things that run the world (the importance and conservation of invertebrates). Conservation Biology 1:344–346.

Wilson, R. J., Z. G. Davies, and C. D. Thomas. 2007. Insects and climate change: processes, patterns and implications for conservation. Pages 245–279 Insect Conservation Biology. Proceedings of the Royal Entomological Society’s 22nd Symposium. CAB International Publishing.

Yu, G., D. K. Smith, H. Zhu, Y. Guan, and T. T. Lam. 2017. ggtree: an R package for visualization and annotation of phylogenetic trees with their covariates and other associated data. Methods in Ecology and Evolution 8:28–36.

Zhang, J., Q. Cong, J. Shen, P. A. Opler, and N. V Grishin. 2019. Genomics of a complete butterfly continent. BioRxiv:829887.

