## Appendix S1 for "Assessing risk for butterflies in the context of climate change, demographic uncertainty, and heterogenous data sources"

**SUPPLEMENTARY MATERIAL FOR:**

### Assessing risk for butterflies in the context of climate change, demographic uncertainty, and heterogenous data sources

Matthew L. Forister[1], Eliza M. Grames[1], Christopher A. Halsch[1], Kevin J. Burls[2], Cas F. Carroll[1], Katherine L. Bell[1], Joshua P. Jahner[3], Taylor Bradford[1], Jing Zhang[4], Qian Cong[5], Nick V. Grishin[4,6], Jeffrey Glassberg[7,8], Arthur M Shapiro[9], Thomas V. Riecke[10]

**Contents included here:**

Table S1 -- results from models predicting risk among group A species.

Table S2 -- taxonomic key for differences between NABA and Pelham lists.

Figure S1 -- perspectives on sampling date as a factor in models of NABA data.

Figure S2 -- examples of land use and climate departure maps.

Figure S3 -- overview of variable transformations.

Figure S4 -- comparison of A and B group species.

Figure S5 -- projected occupancy vs geometric population growth rates.

Figure S6 -- correlation matrix for primary variables.

Figure S7 - S9 -- risk values for all species not shown in Figure 3.

**Other supplementary materials:**

*https://elizagrames.shinyapps.io/butterflyRisk/*

Interactive tool for exploring the impact of different weighting schemes (among contributing variables) on the risk index; also available are species-specific plots and maps similar to main Figures 3 - 5 that can be filtered by state or taxonomic family.

**Table S1.** Coefficients and other results from a Bayesian linear model predicting the composite risk index among A group species using seven predictor variables including climate, land use and natural history. Values shown in the table below are the standardized beta coefficients, upper and lower 95% credible intervals (CI), and probabilities of effect (the fraction of the posterior probability distributions above or below zero, depending on the sign of the coefficient). The variance explained (as the square of the correlation between observed and predicted values) for the whole model was 0.087.

| Variable | Probability | Coefficient | Lower 95% CI | Upper 95% CI |
|---|---|---|---|---|
| Geographic range | 0.58 | -0.0016 | -0.019 | 0.015 |
| Development | 0.52 | -0.00046 | -0.015 | 0.014 |
| Climate departure | 0.92 | -0.013 | -0.032 | 0.0056 |
| Precipitation | 0.98 | -0.021 | -0.040 | -0.0014 |
| Voltinism | 0.51 | -0.00021 | -0.017 | 0.017 |
| Wingspan | 0.99 | -0.019 | -0.033 | -0.0054 |
| Host range | 0.74 | 0.0050 | -0.011 | 0.020 |

**Table S2.** Taxonomic key for differences between the North American Butterfly Association names (NABA, 2018, Checklist of North American Butterflies Occurring North of Mexico, Edition 2.4) and names used in Pelham (Pelham, 2022, A Catalogue of the Butterflies of the United States and Canada). Species names flagged with an asterisk in Figure 2 and Appendix S1: Figures S7 - S9 are shown here in the NABA columns, with the corresponding names in the Pelham columns.

| NABA checklist | Pelham 2022 | NABA checklist | Pelham 2022 |
|---|---|---|---|
| *Achalarus casica* | *Thorybes casica* | *Nymphalis vaualbum* | *Nymphalis l-album* |
| *Adelpha bredowii* | *Adelpha californica* | *Oarisma edwardsii* | *Copaeodes edwardsii* |
| *Agraulis vanillae* | *Dione incarnata* | *Oeneis taygete* | *Oeneis bore taygete* |
| *Amblyscirtes elissa* | *Amblyscirtes arizonae* | *Papilio canadensis* | *Pterourus canadensis* |
| *Atrytonopsis edwardsii* | *Atrytonopsis edwardsi* | *Papilio cresphontes* | *Heraclides cresphontes* |
| *Autochton cellus* | *Telegonus cellus* | *Papilio eurymedon* | *Pterourus eurymedon* |
| *Boloria montinus* | *Boloria chariclea* | *Papilio glaucus* | *Pterourus canadensis* |
| *Boloria napaea* | *Boloria alaskensis* | *Papilio multicaudata* | *Pterourus multicaudata* |
| *Brephidium exile* | *Brephidium exilis* | *Papilio rutulus* | *Pterourus rutulus* |
| *Carterocephalus palaemon* | *Carterocephalus skada* | *Phaeostrymon alcestis* | *Satyrium alcestis* |
| *Chioides catillus* | *Chioides albofasciatus* | *Pholisora mejicana* | *Pholisora mejicanus* |
| *Chiomara asychis* | *Chiomara georgina* | *Phyciodes campestris* | *Phyciodes pulchella* |
| *Colias cesonia* | *Zerene cesonia* | *Phyciodes selenis* | *Phyciodes cocyta selenis* |
| *Colias eurydice* | *Zerene eurydice* | *Phyciodes texana* | *Anthanassa texana* |
| *Copaeodes aurantiacus* | *Copaeodes aurantiaca* | *Phyciodes vesta* | *Phyciodes graphica* |
| *Copaeodes minimus* | *Copaeodes minima* | *Pieris napi* | *Pieris oleracea* |
| *Dymasia dymas* | *Microtia dymas* | *Piruna cingo* | *Piruna aea* |
| *Emesis ares* | *Apodemia ares* | *Plebejus acmon* | *Icaricia acmon* |
| *Emesis zela* | *Apodemia zela* | *Plebejus emigdionis* | *Plebulina emigdionis* |
| *Erebia theano* | *Erebia pawloskii* | *Plebejus icarioides* | *Icaricia icarioides* |
| *Eurema boisduvaliana* | *Abaeis boisduvaliana* | *Plebejus lupini* | *Icaricia lupini* |
| *Eurema dina* | *Pyrisitia dina* | *Plebejus neurona* | *Icaricia neurona* |
| *Eurema lisa* | *Pyrisitia lisa* | *Plebejus saepiolus* | *Icaricia saepiolus* |
| *Eurema mexicana* | *Abaeis mexicana* | *Plebejus shasta* | *Icaricia shasta* |
| *Eurema nicippe* | *Abaeis nicippe* | *Poanes hobomok* | *Lon hobomok* |
| *Eurema nise* | *Pyrisitia nise* | *Poanes melane* | *Lon melane* |
| *Eurema proterpia* | *Pyrisitia proterpia* | *Poanes taxiles* | *Lon taxiles* |
| *Everes amyntula* | *Cupido amyntula* | *Pontia beckerii* | *Pontieuchloia beckerii* |
| *Everes comyntas* | *Cupido comyntas* | *Pontia sisymbrii* | *Sisymbria sisymbrii* |
| *Ganyra howarthii* | *Ganyra howarthi* | *Pyrgus albescens* | *Burnsius albescens* |
| *Hemiargus isola* | *Echinargus isola* | *Pyrgus communis* | *Burnsius communis* |
| *Junonia genoveva* | *Junonia neildi* | *Pyrgus oileus* | *Burnsius oileus (Lin* |

| | | | |
|---|---|---|---|
| *Lycaeides idas* | *Plebejus idas* | *Pyrgus philetas* | *Burnsius philetas* |
| *Lycaeides melissa* | *Plebejus melissa* | *Pyrrhopyge araxes* | *Apyrrothrix araxes* |
| *Lycaena arota* | *Tharsalea arota* | *Satyrodes eurydice* | *Lethe eurydice* |
| *Lycaena dione* | *Tharsalea dione* | *Speyeria adiaste* | *Argynnis adiaste* |
| *Lycaena editha* | *Tharsalea editha* | *Speyeria aphrodite* | *Argynnis aphrodite* |
| *Lycaena gorgon* | *Tharsalea gorgon* | *Speyeria atlantis* | *Argynnis atlantis* |
| *Lycaena helloides* | *Tharsalea helloides* | *Speyeria callippe* | *Argynnis callippe* |
| *Lycaena hermes* | *Tharsalea hermes* | *Speyeria coronis* | *Argynnis coronis* |
| *Lycaena heteronea* | *Tharsalea heteronea* | *Speyeria cybele* | *Argynnis cybele* |
| *Lycaena hyllus* | *Tharsalea hyllus* | *Speyeria edwardsii* | *Argynnis edwardsii* |
| *Lycaena mariposa* | *Tharsalea mariposa* | *Speyeria egleis* | *Argynnis egleis* |
| *Lycaena nivalis* | *Tharsalea nivalis* | *Speyeria hydaspe* | *Argynnis hydaspe* |
| *Lycaena rubidus* | *Tharsalea rubidus* | *Speyeria idalia* | *Argynnis idalia* |
| *Lycaena xanthoides* | *Tharsalea xanthoides* | *Speyeria mormonia* | *Argynnis mormonia* |
| *Megisto rubricata* | *Cissia rubricata* | *Speyeria nokomis* | *Argynnis nokomis* |
| *Neominois ridingsii* | *Oeneis ridingsii* | *Speyeria zerene* | *Argynnis zerene* |
| *Neophasia terlootii* | *Neophasia terlooii* | *Texola elada* | *Microtia elada* |
| *Nymphalis milberti* | *Aglais milberti* | *Thorybes mexicanus* | *Thorybes nevada* |

**Figure S1**. Consideration of sampling date as a predictor for abundance from the NABA data. As described in the main text, our primary model for the NABA data included (among other variables) random effects of site and year, but did not incorporate the date of sampling. The date of sampling does indeed vary through time at each site, as in panel (a), where sampling dates are shown for the sites used in our core NABA model. Each line in the figure is a different site; 2 sites out of 44 were excluded from this graph because they had earlier sampling times in the spring and were not easily visualized on the same graph.

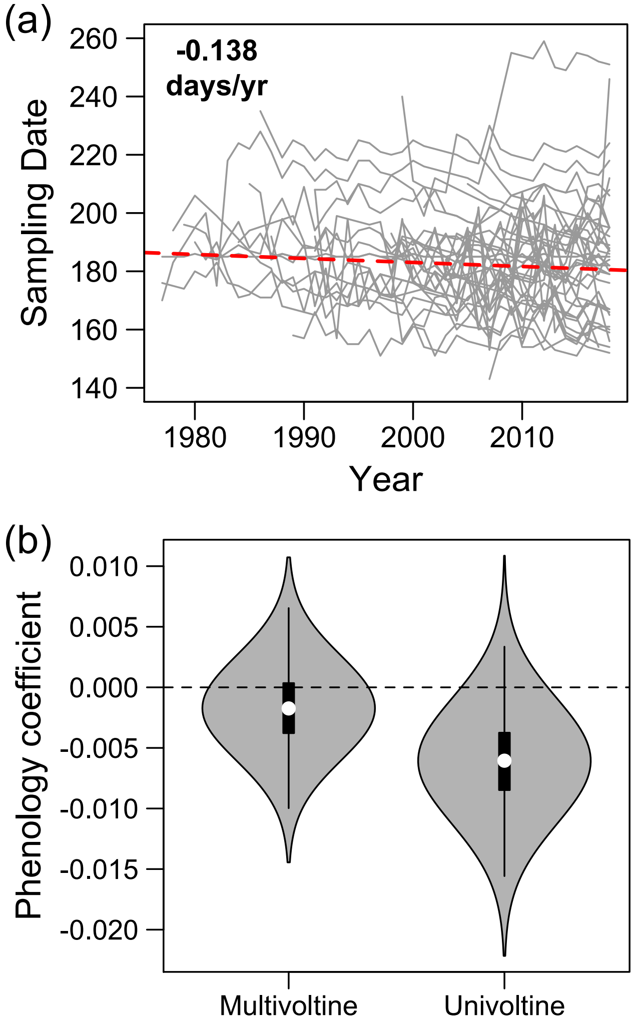


The slope shown in the upper left of panel (a), and the dotted red line are the result of a mixed-effects model executed with the `lme4`[1] package in R. We modeled survey date (*d*) at each site (*i*) in each year (*t*) as a function of a fixed, continuous effect of year with random site-specific intercepts to account for variation in survey timing among sites. Thus, our model was,

$$d_{it} \sim \text{Normal}(\alpha + \beta \times t + \varepsilon_i, \sigma_d^2)$$
$$\varepsilon_i \sim Normal(0, \sigma_\varepsilon^2)$$

We found that the date of sampling across sites has been shifting to an earlier date, at a rate of 0.138 days per year (t value = -5.09), or slightly more than 1 day per decade.

In order to encompass that sampling date variation in analysis of the NABA data, we built a second hierarchical Bayesian model to ask if the inclusion of sampling date would affect our results or produce additional insights. The details of this model are not shown here, but in essence it was identical to the NABA model described in the main text (see Methods section titled **Variable creation part 1**), with the addition of sampling date as a continuous variable. That new, continuous variable was associated with a beta coefficient estimated at each site and separately within the pool of multivoltine and univoltine species.

We expected the date of the NABA census to have relatively little effect on multivoltine species that are present for a long flight window; on the other hand, we hypothesized that sampling date might have a negative association with univoltine species. The latter effect could be generated by later sampling dates missing the peak of abundance, along with the

[1] Bates, D., Mächler, M., Bolker, B., Walker, S. 2015. Fitting linear mixed-effects models using lme4. Journal of Statistical Software 67:1–48.

complementary possibility that warming temperatures from climate change might be advancing the start of the flight season.

In agreement with these possibilities, our model including the sampling date term estimated a close-to-zero effect for multivoltine species, but a negative effect of sampling date for univoltine species, as shown in violin plots of posterior distributions for those two effects of the "phenology coefficient" in panel (b). The negative effect for univoltine species reflects the effect (as hypothesized) that later season sampling in some years for these species returns predictably lower counts.

Finally, we asked if this new model including sampling date would return results that differed in an important way relative to our core model reported in the main text. For each species at each site, we used both models (the core model and the model reported here) to generate predicted values of abundance during the last year of data. We found that those predicted values between the two model outputs were correlated at $r = 0.9999864$. Thus we conclude that the inclusion of sampling date in models of the NABA data produces meaningful results with respect to differences among uni- and multivoltine species as shown in panel (b), but does not change key model output to the extent that would require replacement of our core model.

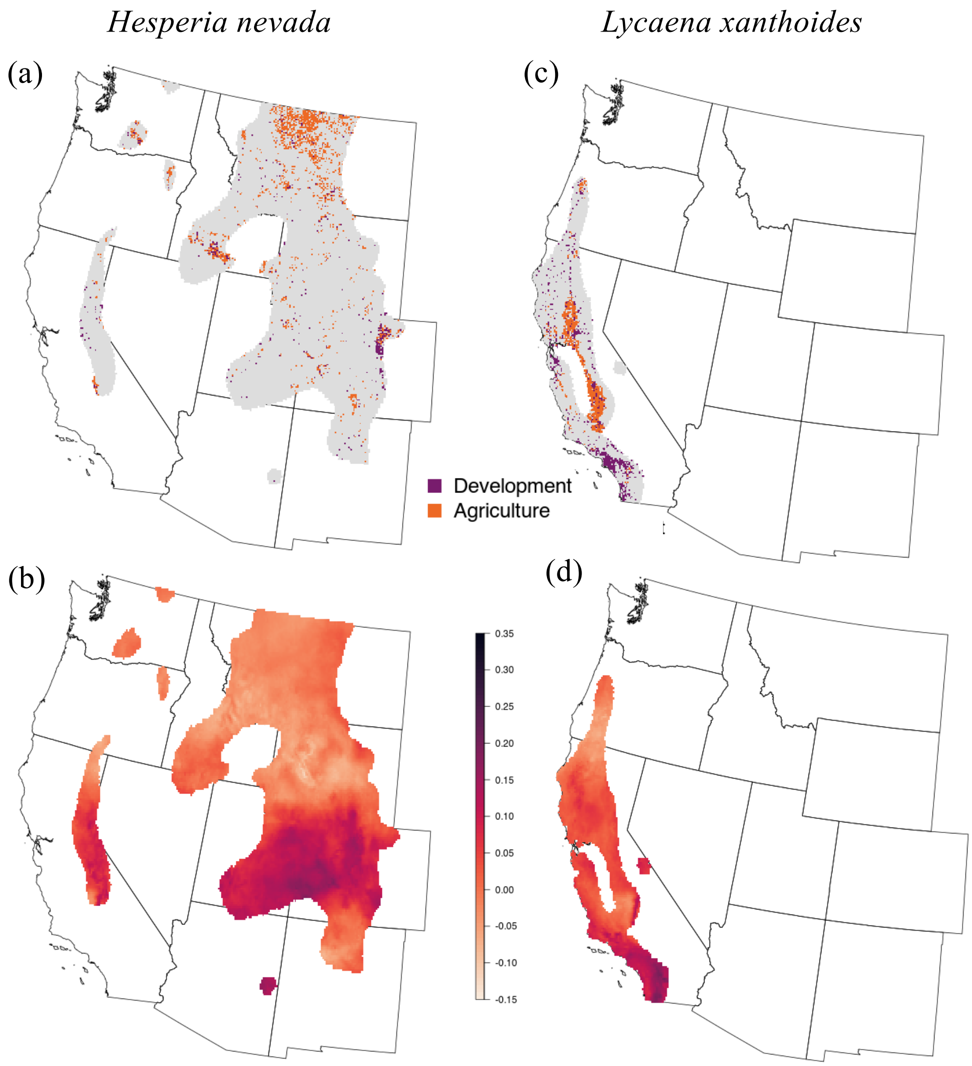


**Figure S2**. Examples of expert range outlines for two species: *Hesperia nevada* in panels (a) and (b), and *Lycaena xanthoides* in panels (c) and (d). In the top two panels, the range is colored by exposure to developed and agricultural lands; in the bottom two panels, the range is colored by multivariate departure from baseline climate conditions. *Lycaena xanthoides*, for example, has a smaller range concentrated in developed parts of California (c), which gives it a higher risk (relative to *H. nevada*) with respect to exposure to land use: this is summarized with a larger risk circle for development in Figure 2. In contrast, the two species have comparable range-wide exposure to climate change (and similar risk circles for that variable in Figure 2).

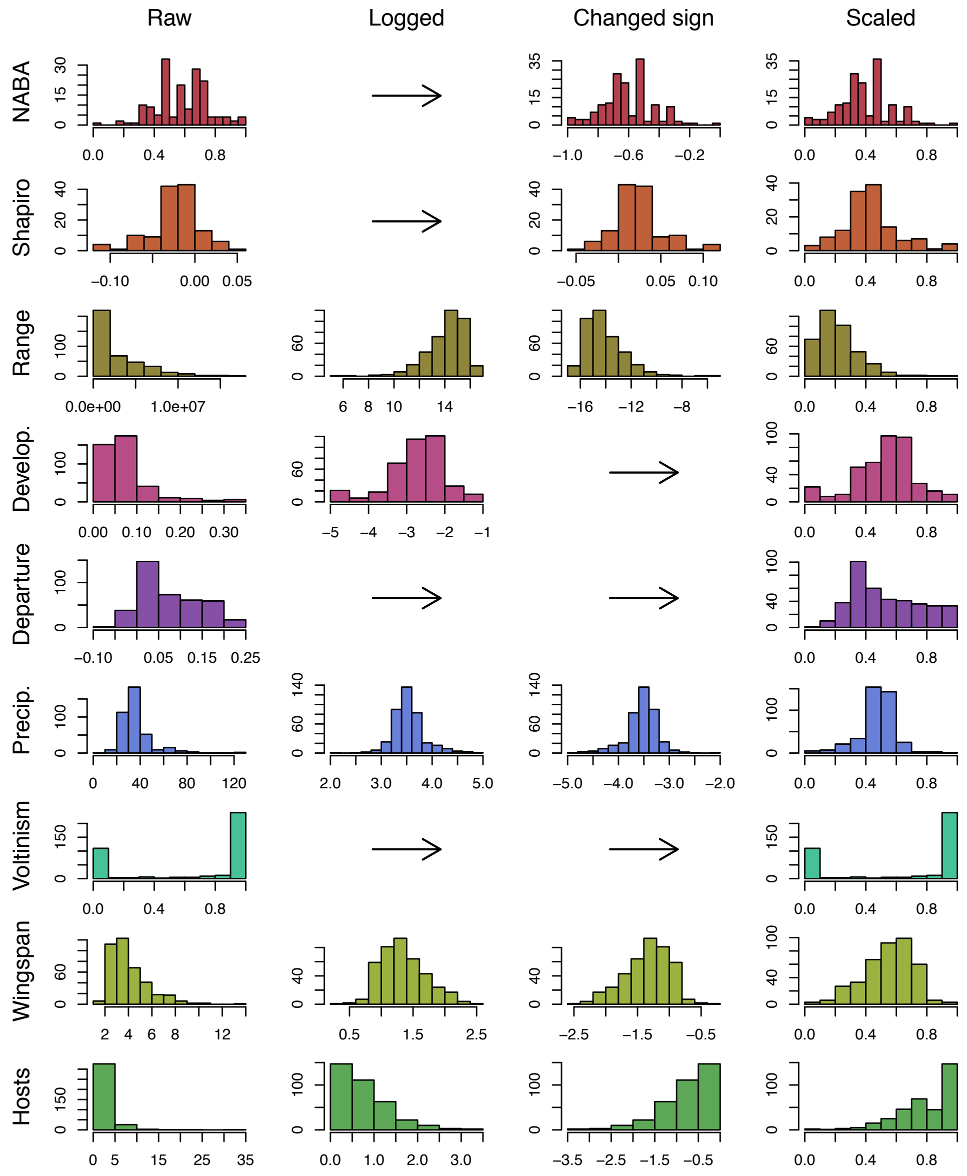


**Figure S3**. Illustration of the process (going from left to right) by which variables were transformed from raw scales into variables bounded between zero and 1, such that higher values correspond to greater risk. For example, NABA values (top row) are probabilities of population persistence (in 50-year simulations), and the signs are changed (positive to negative) so that lower probabilities of persistence are then farther to the right on the x axis; in the last step, the distribution was shifted to the positive by addition of the absolute value of the smallest (most negative) value, and finally divided by the largest value, thus scaling the numbers between 0 and 1. In contrast, the signs were not reversed for development ($4^{th}$ row) because higher values naturally correspond to higher risk (but development values are logged because of high skew, and of course scaled to be between 0 and 1). An arrow indicates a transformation not applied to a given variable. Colors here match colors in Figure 2 and Appendix S1: Figures S7 – S9.

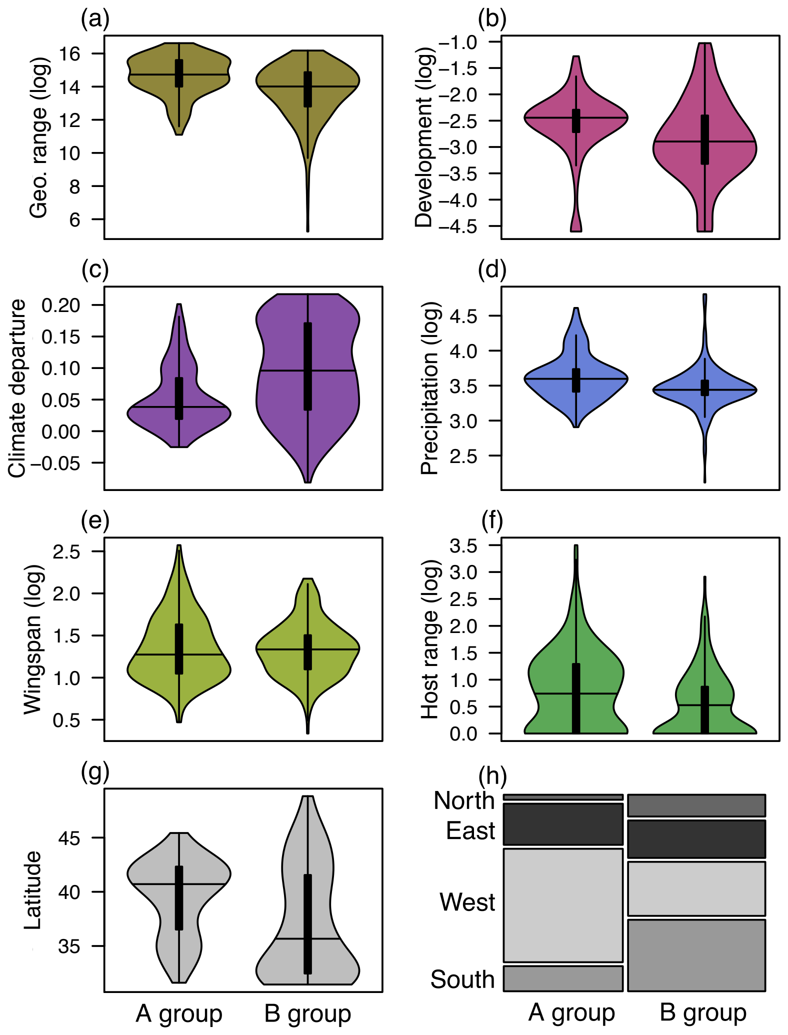


**Figure S4.** Summary of differences between species in the A and B groups. The 184 A group species are those with observational data from either the Shapiro monitoring program or the NABA annual counts; the 212 B group species are not included in those data sources (at least not with sufficient abundance to be used in our primary models). Comparisons in panels (a) through (g) are shown as violin plots with kernel density estimates and horizontal lines marking medians inside rectangles spanning interquartile ranges; vertical lines are upper and lower fences computed as the third quartile plus one and a half times the interquartile range, and the first quartile minus one and a half times the interquartile range, respectively. Colors in panels (a) through (f) match those used in Figure 2 for the same variables. Area-weighted latitudinal midpoints are shown in panel (g), and the mosaic plot in (h) shows the biogeographical breakdown of qualitative range positions for A and B group species (e.g., species with ranges in the South category have a majority of their range south of the US-Mexico border, with only a small presence north of the border in the western US). The latitudinal midpoints (g) were calculated using the rasterToPoints function in the `raster` package v3.5-11[2].

---

[2] Hijmans, R.J., van Etten, J., Mattiuzzi, M., Sumner, M., Greenberg, J.A., Lamingueiro, O.P., Bevan, A., Racine, E.B. & Shortridge, A. (2021). raster: Geographic Data Analysis and Modeling, Version 2.9-23, R package.

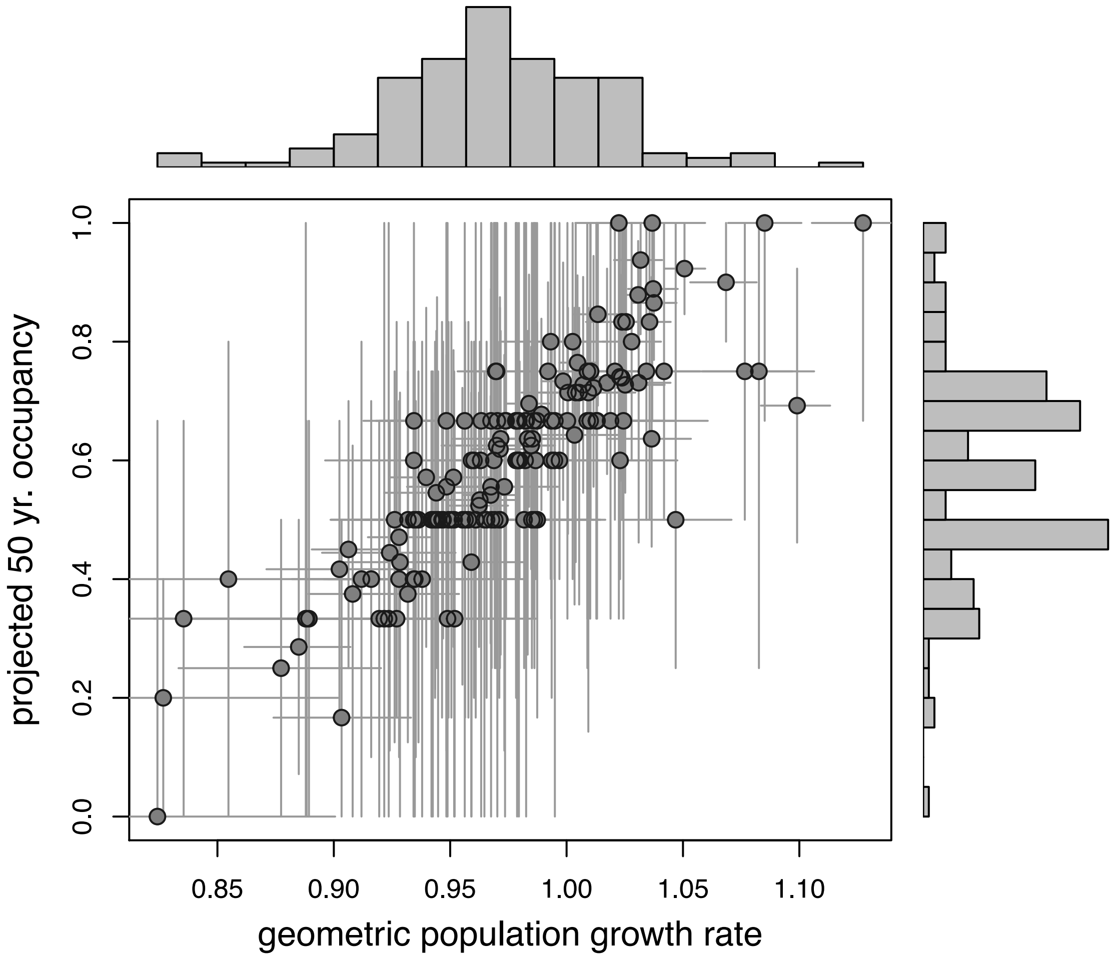


**Figure S5**. Relationship between two variables derived from the NABA community model, in which detection information is shared among species. Points are individual species, with 85% highest density intervals for both axes. The y axis is the projected 50 year occupancy across populations for each species (also referred to in the main text as the probability of population persistence); the x axis is the geometric population growth rate. The latter (growth rate) influences the former (occupancy) in simulations, therefore a positive relationship between these variables is expected (and observed), but we present it here as an illustration of the variation in the two aspects of our model. Marginal histograms show the distribution of among-species variation for values along both axes.

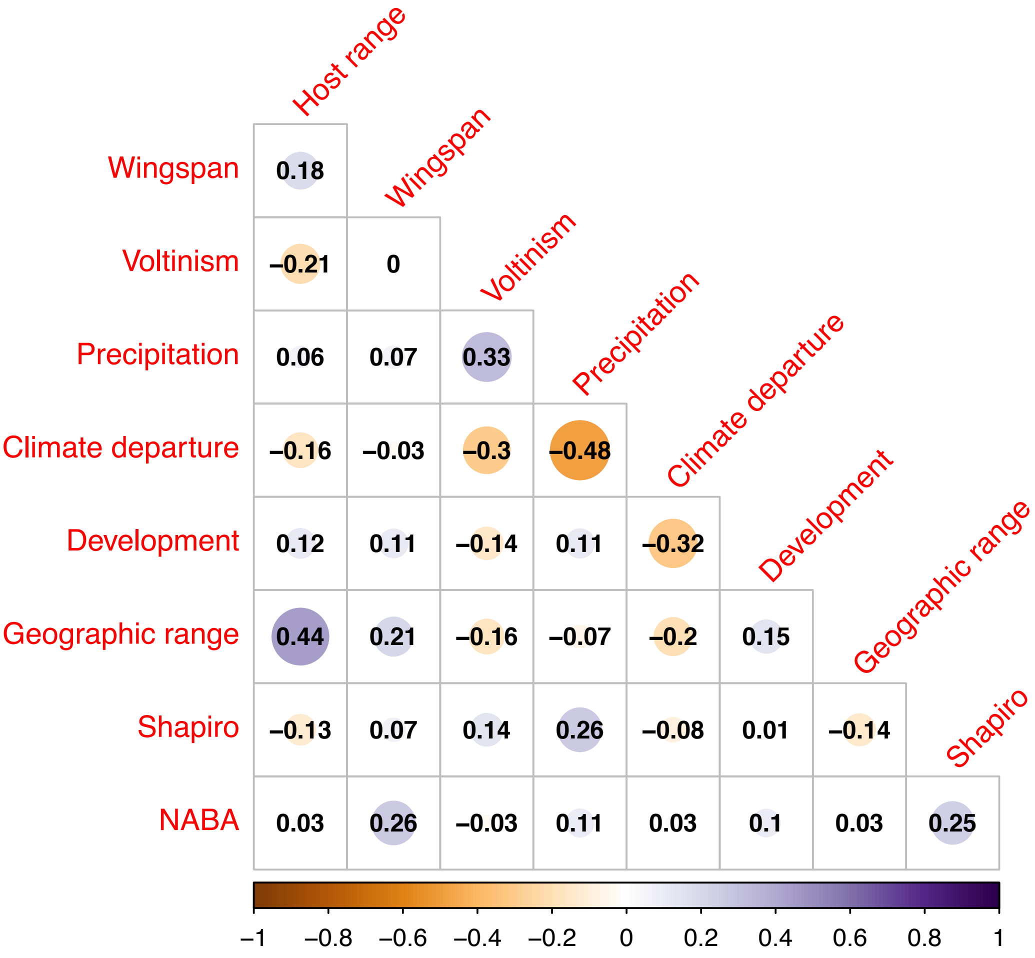


**Figure S6.** Pearson correlation coefficients among all variables shown in main risk plots (Figure 2 and Appendix S1: Figures S7 – S9). Underlying circles and colors are shown for ease of visualizing relative magnitude of values.

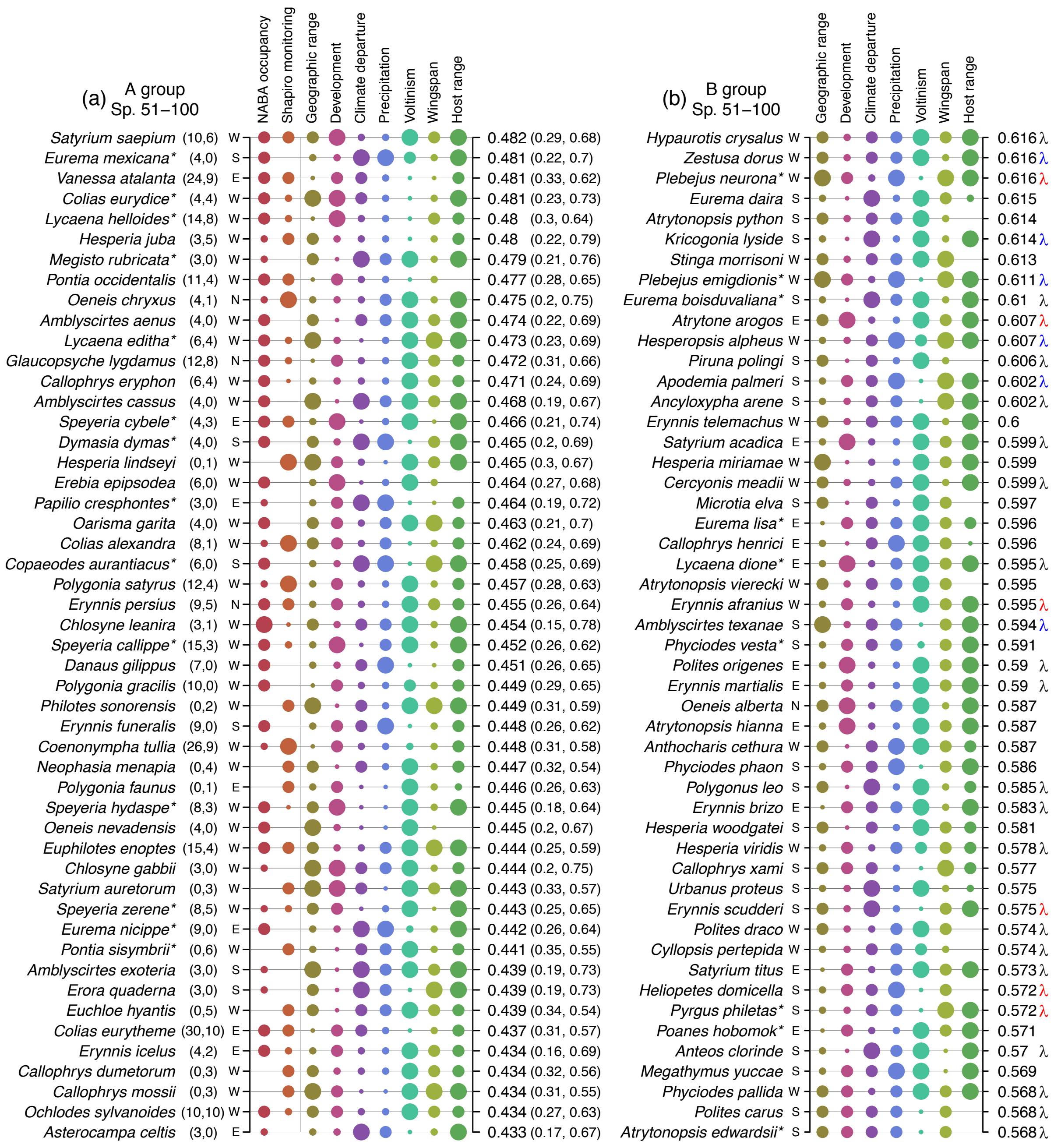


**Figure S7.** The second set of species (51 – 100 for both A and B groups) ranked by risk index values. See Figure 2 for the first set (species 1 – 50), and also see the Figure 2 legend for a full description of the features of the plot shown here.

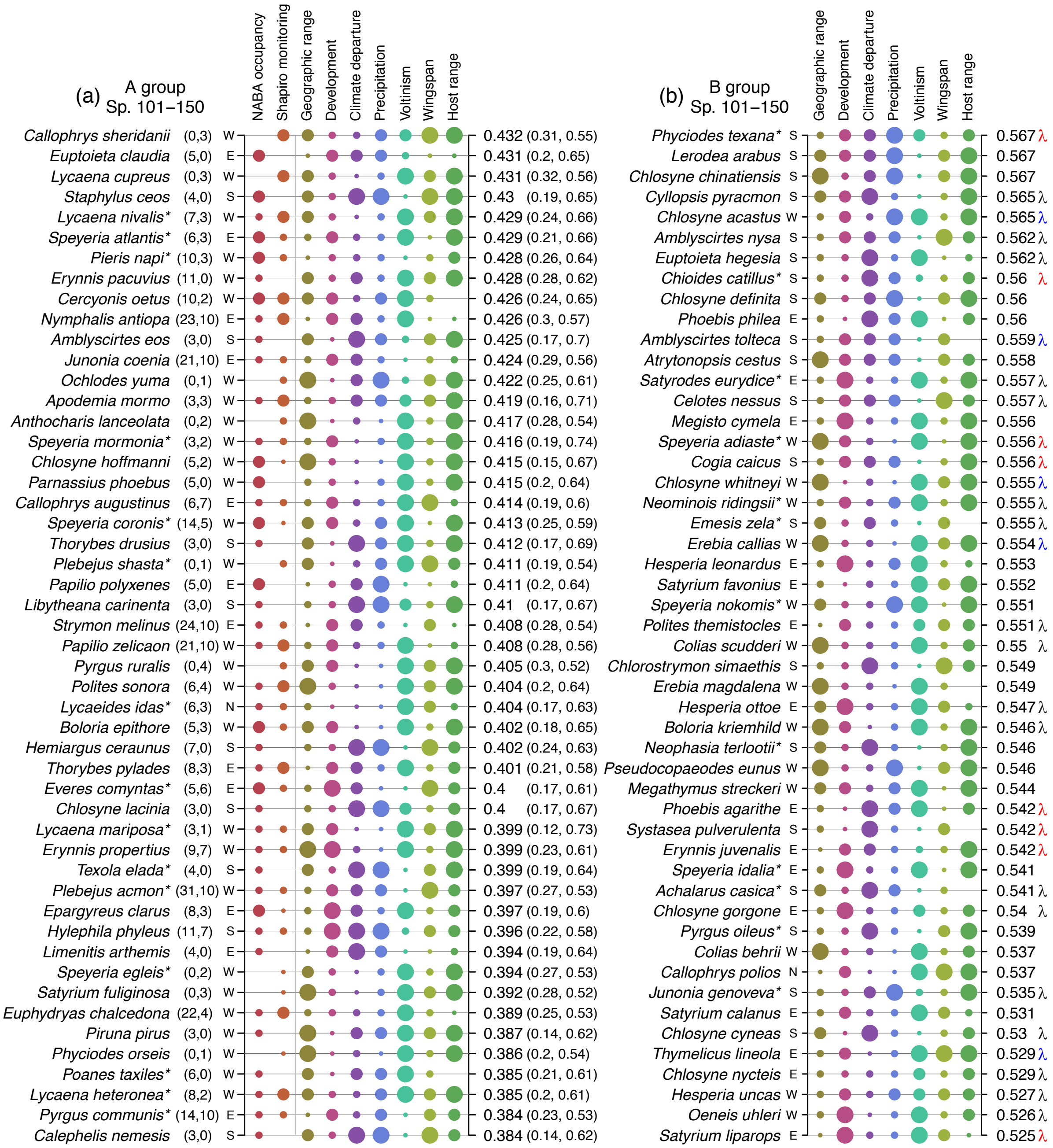


**Figure S8.** The third set of species (101 – 150 for both A and B groups) ranked by risk index values. See Figure 2 legend for a description of all features of the plot.

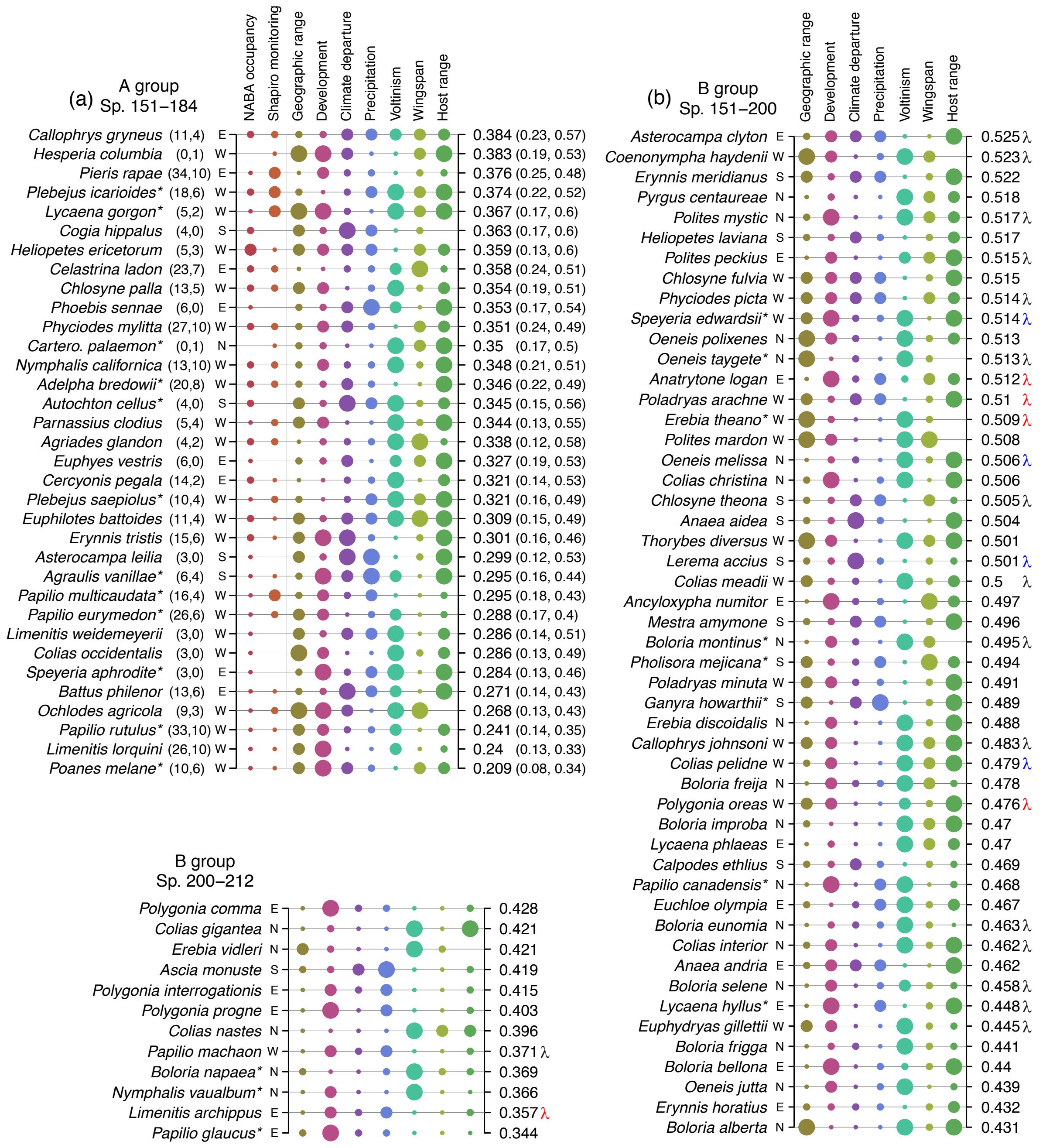


**Figure S9.** The fourth set of species (101 – 151 for the A group, and 151 – 212 for the B group) ranked by risk index values. See Figure 2 legend for a description of all features of the plot, and note here that the last species in the B group (200 – 212) are shown in the lower left of the plot.
